# The single nucleotide polymorphism rs1053230 modulates kynurenine 3-monooxygenase stability and is associated with cognitive and mood phenotypes

**DOI:** 10.64898/2026.05.19.726163

**Authors:** Mary E W Collier, Georgia Ceeney, Joshua Chiappelli, Sathyasaikumar Korrapati, Thong Huy Cao, Paulene Quinn, Junfeng Ma, Aashti Shauriq, Nicolas Sylvius, Edward J Hollox, Donald J L Jones, Andrew Hudson, Elliot L Hong, Nigel Scrutton, Robert Schwarcz, Flaviano Giorgini

## Abstract

**Background:** The single nucleotide polymorphism (SNP) rs1053230 within the kynurenine 3-monooxygenase (*KMO*) gene encodes either an arginine (<u>C</u>GC) or cysteine (<u>T</u>GC) at amino acid residue 452. The rs1053230 genotype is associated with alterations in KMO expression and activity, and impaired cognition. Additionally, KMO intronic SNP rs2275163 is associated with schizophrenia endophenotypes. However, the direct functional consequences of these SNPs on KMO function have never been investigated.

**Methods:** Here we performed the first *in vitro* cell-based examination of the rs1053230 genotype on KMO expression, activity, cellular localisation and KMO-protein interactions, as well as examination of the effects of rs1053230 on schizophrenia-relevant clinical measures. We also examined the effects of rs2275163 genotype on *KMO* pre-mRNA stability and alternative splicing.

**Results:** HEK293T cells expressing KMO-Arg452 or KMO-Cys452 with a red fluorescent protein (RFP) tag produced equivalent levels of KMO mRNA, protein and enzymatic activity, and localised to mitochondria to the same extent. However, cycloheximide-mediated inhibition of protein translation revealed a striking reduction in protein stability of KMO-Arg452-RFP. KMO-RFP-trap pull-down followed by tandem liquid-chromatography-mass spectrometry (LC-MS/MS) identified dramatic differences in protein partners between KMO variants. Indeed, gene ontology-term enrichment analysis revealed that terms associated with synaptic function were more highly enriched amongst KMO-Cys452 interacting proteins. rs1053230 genotype was found to associate with chronic, trait-like depressive mood symptoms in patients. rs2275163 genotype had no effect on *KMO* pre-mRNA.

**Conclusions:** Differences in protein stability and protein-protein interactions may underlie the mechanisms by which the KMO rs1053230 genotype influences neuronal function, leading to cognitive differences in psychiatric conditions.

## Introduction

The kynurenine pathway (KP) is the major route of tryptophan degradation in mammals and accounts for 95 % of the catabolism of this amino acid. Kynurenine 3-monooxygenase (KMO) is a key enzyme in the KP that converts L-kynurenine (L-KYN), a metabolite of tryptophan degradation, to 3-hydroxykynurenine (3-HK). An alternative route for L-KYN degradation is the conversion of L-KYN to kynurenic acid (KYNA) by kynurenine aminotransferase II (KATII). Under physiological conditions, the KMO branch of the KP is the main route for L-KYN metabolism, resulting in the production of 3-HK which is further metabolised to quinolinic acid (QUIN). However, a reduction in KMO activity can cause a shift of the KP towards the KATII branch, resulting in increased levels of KYNA. Several KP metabolites are neuroactive including KYNA [1], 3-HK [2], and QUIN [3] and have been implicated in both psychiatric and neurodegenerative conditions (reviewed in [4] and [5]). In particular, several studies have shown a strong link between increased levels of KYNA and schizophrenia [6,7,8].

Schizophrenia is a severe disease that affects approximately 1 % of the population [9] and presents as positive symptoms such as hallucinations and delusions, negative symptoms including loss of emotion and social withdrawal, and cognitive symptoms such as problems with memory and learning. In addition to more well-known hypotheses of schizophrenia such as the dopamine hypothesis [10], the KP has also been implicated in the pathophysiology of the disease [11]. This has been linked to reduced *KMO* mRNA expression in the frontal cortex of patients [12] and corresponding increases in KYNA levels in the prefrontal cortex [6,7] and cerebrospinal fluid (CSF) [8,13,14]. Increased levels of KYNA in the central nervous system (CNS) have been implicated particularly in the cognitive deficits observed in schizophrenia and may be due to the inhibitory action of this metabolite at N-methyl-D-aspartate receptors (NMDAR) [1] and alpha7 nicotinic acetylcholine receptors (α7nAChR) [15]. In support of this hypothesis, elevated brain levels of KYNA have been consistently shown to result in impairments in memory and learning in rodent models [16,17,18]. Thus, reduced KMO activity, resulting in a shift in the KP towards the KYNA branch, may lead to cognitive dysfunction.

KMO is an outer mitochondrial membrane protein [19] which is expressed in the CNS in microglia [20] and neurons [21]. Two single nucleotide polymorphisms (SNPs) within the *KMO* gene have been previously associated with schizophrenia, the coding SNP rs1053230, located in exon 15 of the KMO gene [22], and the intronic SNP rs2275163 within intron 9 [12]. The non-synonymous coding SNP rs1053230 results in an amino acid substitution; the C genotype encodes an arginine residue at amino acid 452 (<u>C</u>GC) (the major variant), whereas the minor T allele (minor allele frequency = 0.29) encodes a cysteine residue at this position (<u>T</u>GC). Previous studies have associated the Arg452 variant with reduced *KMO* expression in lymphoblastoid cells and hippocampal biopsies from patients with epilepsy [22], increased levels of KYNA in the CSF of patients with bipolar disorder [22] and reduced cognitive function [23]. Together these studies suggest that the rs1053230 genotype affects KMO function and may thus have a role in exacerbating symptoms in individuals with schizophrenia. However, the differences in KMO expression and activity detected in these studies were observed in patient samples which are complex and where other factors may play a role. Therefore, this study directly examined any potential functional differences between the Arg452 and Cys452 variants (e.g. enzymatic activity or other non-enzymatic properties) and any consequent downstream cellular perturbations using a cell-based *in vitro* model. Since the intronic KMO SNP rs2275163 has been associated with schizophrenia endophenotypes [12], we also examined in detail the molecular mechanisms by which this SNP may alter *KMO* pre-mRNA stability and alternative splicing using cell-based *in vitro* models.

## Methods and Materials

### Cell culture and transient transfection

HEK293T cells and the microglial cell line C20 [24] were cultured in DMEM supplemented with 10 % (v/v) foetal calf serum (FCS) under 5 % CO_2_ at 37°C. Prior to transfection, cells were seeded into 6-well plates at a density of 2.5 × 10^5^ cells per well in 2 ml of DMEM containing 10 % FCS (v/v) and incubated overnight at 37°C. Cells were transfected with 2 µg of endotoxin-free plasmid DNA using 2 µl of Lipofectamine 3000 (Thermo Fisher Scientific, Loughborough, UK). Following 6 h transfection, the media was replaced with fresh DMEM containing 10 % FCS and incubated for 24-48 h at 37°C.

The THP-1 monocytic cell line was cultured in RPMI 1640 supplemented with 10 % (v/v) FCS under 5 % CO_2_ at 37°C. Cells were maintained at a density of 1.5 to 8×10^5^/ml. Prior to experiments, THP-1 cells were differentiated into a monocytic phenotype by incubation with 50 nM of 1α,25-dihydroxyvitamin D3 (Sigma-Aldrich, Gillingham, UK) for 72 h [25].

### Plasmids and site-directed mutagenesis

The pcDNA3.1-KMO-RFP plasmid encoding full length KMO with a C-terminal red fluorescent protein (RFP) tag has been described before [26] was verified by Sanger sequencing, which revealed that this vector encoded KMO with the C genotype of rs1053230 resulting in an arginine at amino acid residue 452 of KMO. Site-directed mutagenesis was used to convert <u>C</u>GC to <u>T</u>GC in order to encode a cysteine residue at position 452 of KMO using the Q5 site directed mutagenesis kit (New England Biolabs (NEB), Hitchin, UK). The KMO-Cys452-RFP insert was then sub-cloned into the empty pcDNA3.1 vector and the sequence confirmed by Sanger sequencing. A pcDNA3.1-KMO-RFP construct encoding truncated KMO (tKMO) that lacks the C-terminal amino acids 420-486 and amino acids 367-379, was examined alongside as a negative control for KMO enzymatic activity and cellular localisation [26].

### RNA isolation and RT-qPCR

Cells were lysed in Trizol (0.5 ml) and RNA isolated using chloroform/isopropanol extraction. RNA was treated with TURBO DNase (Thermo Fisher Scientific) for 30 min at 37°C. RNA (1 µg) was then incubated with gDNA Wipeout buffer (Qiagen, Manchester, UK) at 42°C for 2 min and used for reverse transcription with the QuantiTect RT kit (Qiagen) at 42°C for 15 min followed by inactivation at 95°C for 3 min. cDNA (1 µl) was used for qPCR with the QuantiTect SYBR Green qPCR kit (Qiagen) and the following primers (0.3 µM final concentration): *KMO* forward 5’-GAATGCGGGCTTTGAAGAC-3’, *KMO* reverse 5’-ACAGGAAGACACAAACTAAGGT-3’, *Rplp0* forward 5’-GCTGCTGCCCGTGCTGGTG-3’, *Rplp0* reverse 5’-TGGTGCCCCTGGAGATTTTAGTGG-3’. Samples were run in three technical replicates using the following program on a Roche LightCycler 480 instrument (Roche, Welwyn Garden City, UK): 95°C for 15 min followed by 40 cycles of 94°C for 15 s, 60°C for 60 s, and 72°C for 30 s. The delta-delta Ct (ΔΔCt) method [27] was used to calculate changes in *KMO* expression compared to the house-keeping gene *Rplp0*. The efficiency of the *KMO* and *Rplp0* primers was calculated as 108 % and 97 % respectively.

### Digital droplet PCR (ddPCR) to examine the effect of rs2275163 genotype on allelic imbalance in KMO mRNA expression

Vitamin D3-differentiated THP-1 cells were treated with lipopolysaccharide (LPS) (0.1 µg/ml) from *Escherichia coli* O111:B4 (Sigma) for 24 h to induce the expression of *KMO*. Cells were then incubated with actinomycin D (1-10 µg/ml) for 1 or 4 h to inhibit RNA synthesis. Cells were lysed in Trizol and RNA was isolated and treated with TURBO DNase as described above. 1 µg of RNA per sample was used for cDNA synthesis using the QuantiTect RT kit and a KMO intron 9/10-specific reverse primer (5’-TTTCCCACGAGTGGAATATGG-3’) (0.7 µM) at 42°C for 30 min and inactivated at 95°C for 3 min. ddPCR was carried out using 2 µl of cDNA (or 2 µl of no reverse transcriptase (RT) reactions as controls), ddPCR Supermix for probes (no dUTP) (Bio-Rad Laboratories Ltd, Watford, UK) and the following primers (900 nM) and MGB rs2275163-C/T-specific probes (250 nM) which were designed and synthesised by Integrated DNA Technologies (IDT; Leuven, Belgium):

rs2275163 ddPCR forward primer: 5’-CACCTGATTTCAGTTTGGGTTAAA-3’

rs2275163 ddPCR reverse primer: 5’-TTCCTGCTCACAGCACAATAT-3’

rs2275163-C specific probe: 5’-/56-FAM/CTTAGACTTTTGCTCTAA/3MGB-NFQ-3’

rs2275163-T specific probe: 5’-5HEX/CACTTAAACTTTTGCTC/3MGB-NFQ/-3’

Droplets were generated using the QX200 droplet generator (Bio-Rad). PCR was initiated at 95°C for 10 min followed by 40 cycles of 94°C for 30 s and 58°C for 1 min, and a final 10 min at 98°C. Within 1 h of the PCR finishing, the plates were equilibrated to room temperature, and samples were analysed on the QX200 droplet reader (Bio-Rad) using the QuantaSoft software (Bio-Rad).

### rs2275163 alternative splicing minigene assay

Synthetic DNA sequences corresponding to exon 9, intron 9-10 and exon 10 of the *KMO* gene with either a C or T at rs2275163 were synthesised by IDT and cloned into the mEmerald-C1 vector downstream of green fluorescent protein (GFP) and sequences were confirmed by Sanger sequencing. C20 cells (1 × 10^5^) were seeded into 12-well plates and transfected with mEmerald-C1-rs2275163-C/T plasmids (1 µg) using Lipofectamine 3000 and incubated at 37°C for 24 h. The medium was replaced with macrophage serum free medium (Thermo Fisher Scientific) and cells were incubated for 4 h. Cells were then treated with tumour necrosis factor alpha (TNFα) (50 ng/ml) or hydrogen peroxide (H_2_O_2_) (50 µM) for 4 h. RNA was isolated and treated with TURBO DNase as described above. RNA (50 ng) was converted to cDNA using the LunaScript RT SuperMix kit (NEB). cDNA (2 µl) or no RT reactions (2 µl) were used for PCR with Q5 High Fidelity DNA polymerase (NEB) and forward and reverse primers (0.5 µM each) specific for the mEmerald-C1 plasmid:

Forward GFP Primer: 5′-TCCAAGCTGAGCAAAGAC-3′

Reverse SV40 polyA primer: 5’-GTAACCATTATAAGCTGCAAT-3’

PCR was performed at 98°C for 30 s, followed by 25 cycles of 98°C for 10 s, 62°C for 30 s and 72°C for 1.5 min with a final extension of 72°C for 2 min. PCR products were separated using 1 % (w/v) agarose gel electrophoresis and images were acquired using a G:BOX Chemi XX9 gel imaging system (Syngene, Cambridge, UK). ImageJ was used to analyse the intensity of the bands.

### SDS-PAGE and immunoblot analysis

Cells were lysed in RIPA buffer containing 1 % (v/v) Halt protease inhibitor cocktail (Thermo Fisher Scientific) and incubated on ice for 20 min. Lysates were centrifuged at 10,000 *g* for 5 min at 4°C and protein concentrations were determined using the DC protein assay (Bio-Rad). Proteins (20 µg) were separated on 10-20 % (v/v) Novex Tris-glycine SDS-PAGE gels (Thermo Fisher Scientific) and transferred onto nitrocellulose membranes. Total protein was visualised using the No-Stain Protein Labelling Reagent (Thermo Fisher Scientific).

Membranes were blocked with Tris-buffered saline Tween-20 (TBST) (50 mM Tris-HCl, pH 8, 150 mM NaCl, 1 % (v/v) Tween-20) containing 5 % (w/v) low fat milk for 1 h and then incubated overnight at 4°C with the following primary antibodies diluted in TBST-milk; monoclonal rabbit anti-KMO clone 2493A (R&D Systems, Abingdon, UK) diluted 1:1000 or rat anti-alpha tubulin clone YOL1/34 (Santa Cruz Biotechnology, Heidelberg, Germany) diluted 1:2500. Membranes were washed three times for 5 min each with TBST and then incubated for 1 h at room temperature with an anti-rabbit-HRP conjugated secondary antibody (Vector Labs, Kirtlington, UK) diluted 1 in 5000 or an anti-rat-HRP antibody (Vector Labs) diluted 1 in 10,000. Membranes were washed four times with TBST and developed with Westar Supernova ECL detection reagent (Cyanagen, Bologna, Italy) and imaged using a G:BOX Chemi XX9 gel imaging system. When necessary, membranes were stripped with 200 mM glycine, pH 2.2, 0.1 % (w/v) SDS, 1 % (v/v) Tween-20, washed thoroughly with TBST and re-probed with appropriate primary antibody.

### KMO activity assay

HEK293T cells (3 × 10^6^) were seeded into T75 flasks and incubated at 37°C overnight. Cells were transfected with 15 µg of plasmid DNA to express KMO-Arg/Cys452-RFP, tKMO-RFP or RFP using Lipofectamine 3000 and incubated at 37°C for 48 h. Cells were then washed three times with 10 ml of ice-cold PBS, scraped into 5 ml of ice-cold PBS and centrifuged at 5000 *g* for 5 min at 4°C. Cell pellets were washed in 1 ml of ice-cold PBS and centrifuged again. The supernatant was removed and cell pellets were snap-frozen in liquid nitrogen.

KMO activity was measured by quantifying 3-HK formation from L-kynurenine using HPLC with electrochemical detection as previously described [26]. Protein concentrations were determined by the Lowry method [28].

### Immunofluorescence confocal microscopy

HEK293T cells (0.4 × 10^5^) were seeded into 35 mm glass-base µ-dishes coated with 0.01 % poly-L-lysine. Cells were incubated overnight and then transfected with 0.4 µg of plasmid DNA using 0.4 µl of Lipofectamine 3000. Following transfection, cells were incubated at 37°C for 48 h. Cells were then washed twice with PBS pre-warmed to 37°C and fixed with pre-warmed 4 % (w/v) paraformaldehyde for 20 min at room temperature. Cells were washed three times with PBS and blocked/permeabilised with PBS containing 5 % (v/v) donkey serum (Merck life Science, Gillingham, UK) and 0.2 % (v/v) Triton X-100 for 30 min at room temperature. Cells were incubated at 4°C overnight with a rabbit anti-HtrA2 antibody (R&D Systems) diluted 1:400 in PBS containing 1 % (w/v) bovine serum albumin (BSA) and 0.2 % (v/v) Triton X-100. Cells were then washed three times with PBS containing 0.05% (v/v) Tween-20 (PBST) and stained with Hoechst 33342 (Abcam, Cambridge, UK) diluted 1:2000 in PBS for 2 min. Cells were incubated at room temperature for 1 h with donkey anti-rabbit Northern Lights-493 antibody (R&D Systems) diluted 1:200 in PBS containing 1 % (w/v) BSA and 0.2 % (v/v) Triton X-100. Cells were washed three times with PBST and placed in PBS with 50 % (v/v) glycerol. Cells were visualised using an Olympus FV1000 confocal laser scanning microscope with a 60x objective and images were acquired using the Olympus FluoView FV1000 software. Pearson’s and Mander’s colocalisation coefficients were measured using the JACoP macro in FIJI ImageJ.

### KMO-RFP trap pull-down assays

HEK293T cells (3 × 10^6^) were seeded into T75 flasks and incubated at 37°C overnight. Cells were transfected with either 15 µg of pcDNA3.1-KMO-Arg452-RFP, pcDNA3.1-KMO-Cys452-RFP or pcDNA3.1-RFP using Lipofectamine 3000 (Thermo Fisher Scientific) as described above. Cells were incubated at 37°C for 48 h to allow the expression of KMO-RFP or RFP, and then washed twice with PBS and lysed in 1 ml of ice-cold lysis buffer (10 mM Tris-HCl, pH 7.5, 150 mM NaCl, 0.5 mM EDTA, 0.05 % n-dodecyl β-D-maltoside, 1 % (v/v) Halt protease inhibitor cocktail, 1 % (v/v) Halt phosphatase inhibitor cocktail). Cells were incubated on ice for 30 min and centrifuged at 5000 *g* for 10 min at 4°C. Four aliquots of each lysate sample (250 µl) were diluted 1:1 with dilution buffer (10 mM Tris-HCl, pH 7.5, 150 mM NaCl, 0.5 mM EDTA, 1 % (v/v) Halt protease inhibitor cocktail, 1 % (v/v) Halt phosphatase inhibitor cocktail) and lysates were cleared with Binding Control Agarose (20 µl per aliquot) (Proteintech, Manchester, UK) for 30 min at 4°C with rotation. Agarose beads were removed by centrifugation at 2500 *g* for 5 min at 4°C. The supernatants were transferred to fresh 1.5 ml tubes and incubated with 25 µl of RFP-Trap Agarose (Proteintech) for 1 h at 4°C with rotation. For SDS-PAGE and immunoblot analysis, all four aliquots per sample were combined together and the agarose beads were washed three times with 500 µl of wash buffer (10 mM Tris-HCl, pH 7.5, 150 mM NaCl, 0.5 mM EDTA) by centrifugation at 2500 *g* for 5 min at 4°C. Proteins were eluted from the beads in 30 µl of 2x SDS-PAGE loading buffer and incubated at 95°C for 5 min. The supernatant was retained and proteins were separated by 10 % (w/v) SDS-PAGE. For mass spectrometry analysis, the beads were combined together for each sample and washed twice with wash buffer (500 µl) and three times with 50 mM ammonium bicarbonate (500 µl). The beads were then resuspended in ammonium bicarbonate (150 µl per aliquot).

### Preparation of RFP-trap samples for mass spectrometry

A bicinchoninic acid assay was performed to estimate the amount of protein in each sample. RapiGest (0.063 % v/v) was added to each bead suspension (150 µl) and incubated at 80°C for 1 h. Dithiothreitol (DTT) was added to a final concentration of 5 mM and incubated at 60°C for 30 min. The samples were allowed to cool to room temperature and iodoacetamide (IAA) was added to a final concentration of 10 mM and incubated in the dark for 30 min at room temperature. Trypsin was added to a final molar ratio of 1:25 trypsin:protein and incubated at 37°C overnight. Formic acid (FA) was added to final concentration of 1% (v/v) and the samples were centrifuged at 14000 *g* for 10 min. The supernatants were purified by solid-phase extraction (SPE). An EMPORE C18 column was washed with methanol followed by four washes with 0.1 % FA. The supernatant was loaded onto the column and washed four times with 0.1 % FA. Peptides were then eluted with 0.6 mL 60 % acetonitrile (ACN)/0.1 % FA, followed by 0.6 mL 80 % ACN/0.1 % FA. Solvents were removed by SpeedVac, flash frozen in liquid nitrogen, and then freeze-dried overnight. The peptide pellet was reconstituted in 20 µL of 0.1 % FA and the peptide concentration was calculated using the o-phthalaldehyde assay. Samples for mass spectrometry were prepared to a final peptide concentration of 1 µg/µL in 0.1 % FA and alcohol dehydrogenase enzyme from *S. cerevisiae* was added at a 1:1 molar ratio as an internal standard.

### Mass spectrometry

Peptides were analysed by LC ESI-MS/MS using the Acquity UPLC coupled to a Water Synapt G2S mass spectrometer (Waters Corporation, Manchester, UK). Sample separation was performed using an Acquity UPLC Symmetry C18 trapping column (180 μm × 20 mm, 5 μm), to remove salt and any other impurities, and a HSS T3 analytical column (75 μm × 150 mm, 1.8 μm). Sample separation was achieved using a gradient from 0.1 % FA in HPLC grade water to 0.1 % FA in ACN over 110 min. The injection volume was 2 µL and each sample was analysed in triplicate. Data were analysed using Progenesis QI for Proteomics version 4.2 (Nonlinear Dynamics, Waters Corporation) to identify peptides and proteins using the UniProtKB database. Progenesis QI also provides quality control metrics to give confidence in the data. The protein mass was limited to a maximum of 1000 kDa. Enzymatic digestion was performed using trypsin, allowing up to two missed cleavages.

Carbamidomethylation C was specified as a fixed modification. Variable modifications included deamidation N, oxidation M, and phosphorylation STY. For peptide identification, a minimum of two fragment ions per peptide was required. In addition, at least five fragment ions were required for protein identification, and each protein had to be supported by a minimum of two peptides. A maximum rate of 1 % was set for the false discovery rate (FDR). The top three most abundant peptides for each protein (Hi-3) were employed for protein quantitation. The results were exported to Excel for further analyses.

### Gene ontology-term enrichment analysis and STRING protein interaction network analysis of KMO-protein interactions

Proteins that were shown to interact with RFP were deemed non-specific interactors and were removed from KMO-Arg/Cys452 lists. KMO-interacting proteins with confidence scores ≥10 (based on the number of peptides or unique peptides) were selected for both the KMO-Arg452 or KMO-Cys452 variants and proteins specific for each variant were identified using the Bio Venn analysis software [29]. Proteins unique to each KMO variant and with confidence scores ≥40 were then analysed using gene ontology (GO)-term enrichment analysis using the PANTHER Overrepresentation test (released 07/08/2024) with GO Ontology database (DOI: 10.5281/zenodo.12173881 released 2024-06-17), and Fishers Exact test with an FDR of <0.05. STRING protein interaction network analysis [30] was performed with lists of interacting proteins unique to KMO-Arg452 or KMO-Cys452 and with confidence scores ≥40. STRING interaction scores were set to a high confidence score of 0.7 and all forms of evidence were used.

### Prediction of KMO variant stability using molecular dynamics simulations

After predicting the structures of KMO-Arg452 and KMO-Cys452 using AlphaFold 3, the Amber 24 software package was used to assess the binding stability between cofactor FAD and protein. The PDB structures were loaded, with the ff14SB force field used to describe the protein, and the GAFF force field employed for the parameterization of the compounds. The system was placed in a TIP3P water box with a boundary distance of 10.0 Å from the protein surface, and Na^+^ and Cl^−^ ions were added to neutralize the charges. Energy minimization was performed in two stages. First, constraints were applied to the protein backbone, optimizing only the solvent and ions. Then, the constraints were removed, and a full minimization of the entire system was performed using a combination of the steepest descent method and the conjugate gradient method to ensure system stability. System equilibration was also performed in two steps. First, the system was heated from a low temperature to 300 K under isothermal conditions with the protein backbone constrained.

Subsequently, further equilibration was performed under isobaric conditions to adjust density, maintaining a temperature of 300 K and pressure of 1 bar, gradually preparing the system for production runs. A 300 ns molecular dynamics simulation was conducted under constant temperature (300 K) and pressure (1 bar) conditions, and no constraints, generating trajectories for subsequent analysis. The trajectories were analysed using the CPPTRAJ tool to calculate the root mean square deviation (RMSD), solvent accessible surface area (SASA), root mean square fluctuation (RMSF), and hydrogen bond count.

### Clinical associations of KMO genotype

Genotype for rs1053230 was assessed in a sample of 156 people with a diagnosis of schizophrenia or schizoaffective disorder and 170 people without a psychiatric diagnosis at the time of study. All participants provided written informed consent after being evaluated for capacity to sign consent. Study procedures were approved by the University of Maryland Baltimore Institutional Review Board. Diagnoses were established with the Structured Clinical Interview for the DSM-IV. Participants completed two cognitive measures; the digit symbol coding task, which provides a measure of processing speed [31], and the digit sequencing task, a measure of working memory [32]. Participants were also assessed for overall psychiatric symptoms with the Brief Psychiatric Rating Scale (BPRS) [33] and for negative symptoms with the Brief Negative Symptom Scale [34]. Participants also completed the Maryland Trait and State Depression scales (MTSD) [35], a self-report questionnaire that assesses severity of current symptoms of depression (state depression) and severity of symptoms experienced over the course of adult life (trait depression).

### Statistical analysis

Statistical analyses were carried out using GraphPad Prism version 10.4.2. Specific statistical tests used for individual experiments are described in the figure legends. Data are shown as mean ± standard deviation. A *p*-value < 0.05 was considered significant. SPSS v.29 was used to analyse clinical measures. The rs1053230 genotype was collapsed into two groups, individuals homozygous for the major allele (CC) and minor allele carriers (CT/TT). ANCOVA were performed with diagnosis and rs1053230 genotype as independent factors and age and sex as covariates.

## Results

### rs2275163 genotype has no effect on *KMO* pre-mRNA allelic imbalance

We first sought to determine whether the intronic SNP rs2275163 genotype had any effect on mRNA levels by assessing allelic imbalance of *KMO* pre-mRNA in the THP-1 monocytic cell line, which is heterozygous (C/T) at this SNP. Initially we examined the effect of rs2275163 genotype on allelic imbalance in THP-1 cells following incubation with LPS (0.1 µg/ml) for 24 h to induce *KMO* expression. ddPCR revealed that the rs2275163 C or T genotypes had no effect on the expression of *KMO* pre-mRNA in untreated cells or cells treated with LPS (Figure S1 in Supplemental Data File 1). Furthermore, inhibition of RNA synthesis using actinomycin D (1-10 µg/ml) revealed that both the C and T allele-containing pre-mRNA transcripts were degraded at similar rates in THP-1 cells and therefore had similar stabilities over the 4 h period examined (Figure 1A, Figure S2 in Supplemental Data File 1). Negative controls lacking reverse transcriptase were run for each sample and showed either no copies, or a very low number (<5) of copies per 20 µl, confirming the absence of genomic DNA contamination. These data suggest that rs2275163 genotype has no effect on *KMO* pre-mRNA stability.

**Figure 1.**
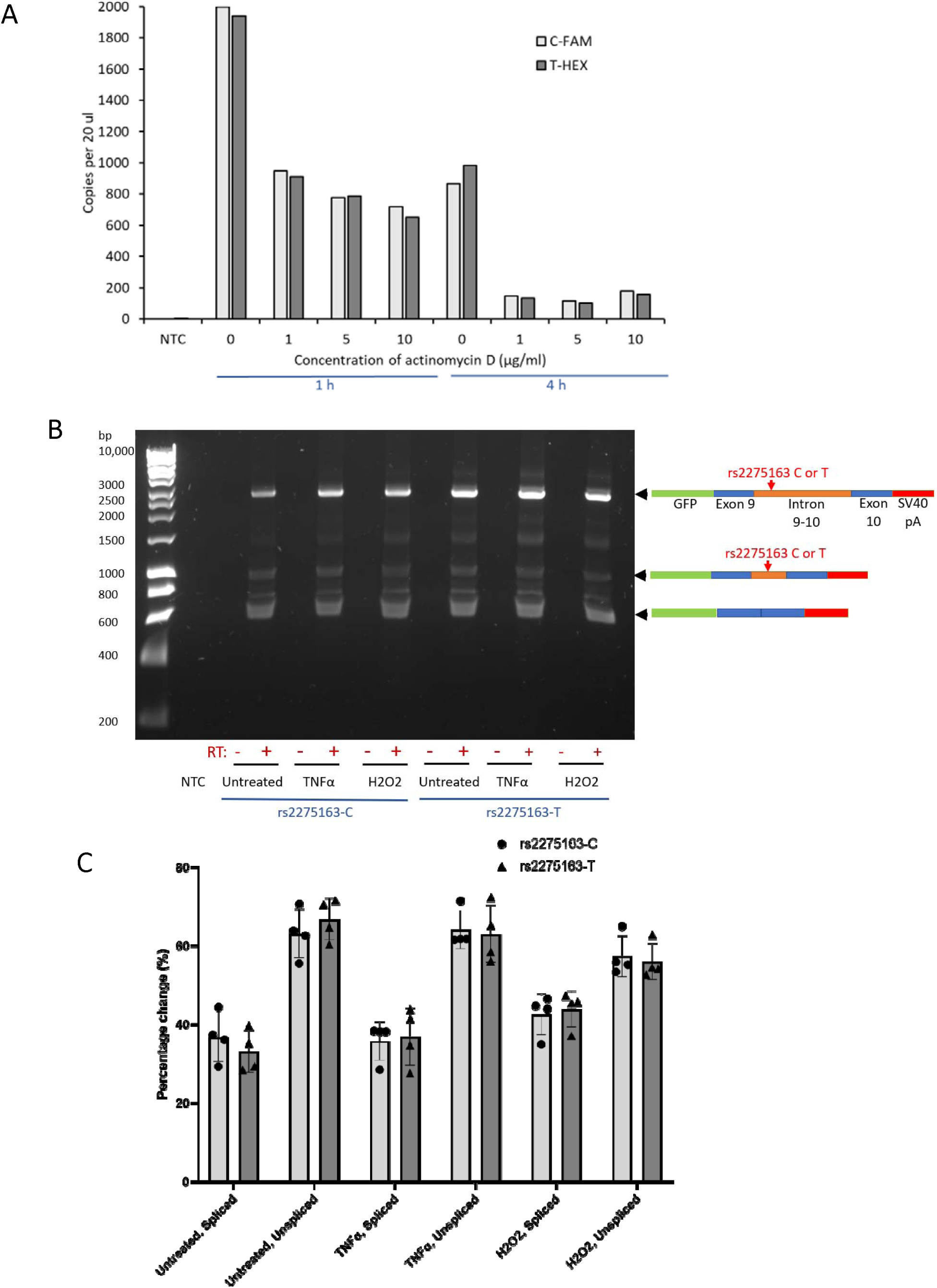
rs2275163 genotype has no effect on *KMO* pre-mRNA stability and alternative splicing. A) THP-1 cells were treated with LPS for 24 h to induce *KMO* expression. Cells were then treated with actinomycin D (1-10 µg/ml) for 1 h or 4 h to inhibit RNA synthesis. RNA was isolated and the effect of rs2275163 genotype on *KMO* pre-mRNA stability was examined using ddPCR (n=1). NTC = no template control. B) C20 cells were transfected with the mEmerald-C1-rs22275163-C/T minigene plasmids and treated with TNFα (50 ng/ml) or H_2_O_2_ (50 µM) for 4 h. PCR was performed with primers specific for the plasmid-derived *KMO* pre-mRNA transcript and PCR products were examined by agarose gel electrophoresis. RT= reverse transcriptase. C) The intensity of the bands was measured using ImageJ and percentage changes in the ratios of spliced and unspliced/partly spliced PCR products were calculated (n=4 ± SD).

### rs2275163 genotype has no effect on alternative splicing of KMO mRNA

Next, we assessed alternative splicing of intron 9-10 of the *KMO* gene containing the rs2275163 C or T genotypes using a minigene expression system [36]. C20 microglial cells were used for this assay because these cells express low levels of KMO [37] and are therefore likely to have the correct complement of factors required for alternative splicing of the *KMO* transcript. Plasmid-specific primers were used to amplify plasmid-derived transcripts to prevent interference from endogenous *KMO* expression [38]. Transcripts containing the C or T genotypes at rs2275163 showed the same patterns of RNA splicing, regardless of rs2275163 genotype. Sanger sequencing was used to confirm the identity of the PCR product. A PCR product of 2462 bp was observed corresponding to unspliced *KMO* exon 9 (122 bp), intron 9-10 (1887 bp) and exon 10 (148 bp), including the GFP and SV40 polyA sequences (305 bp) (Figure 1B). The fully spliced PCR product of 575 bp corresponding to *KMO* exon 9 and exon 10 (including 305 bp of GFP and polyA sequence) was also observed, indicating that the *KMO* minigene mRNA construct expressed from the plasmid was successfully processed in C20 cells (Figure 1B). Furthermore, a partially spliced product of 963 bp corresponding to exon 9, the first 388 bp of intron 9-10 and exon 10 was detected (Figure 1B). This partially spliced PCR product was shown to be spliced at a putative cryptic donor splicing site within intron 9-10 using the Alternative Splice Site Predictor (ASSP) [39]. Densitometry analysis confirmed there were no significant differences between the ratios of spliced and partially spliced or unspliced PCR products in C20 cells, even after stimulation of cells with TNFα or H_2_O_2_ (Figure 1C). Since rs2275163 genotype had no observable effect on *KMO* pre-mRNA stability or alternative splicing, the study then focused only on the KMO coding SNP rs1053230.

### KMO-Arg452 and KMO-Cys452 mRNA and protein expression levels and KMO enzymatic activity are not significantly different in HEK293T cells

HEK293T cells were transfected to express KMO with the C and T genotypes at SNP rs1053230 to express KMO-Arg452 and KMO-Cys452 respectively. Analysis of *KMO*-Arg452 and Cys452 mRNA expression in HEK293T cells 24 h post transfection revealed there were no significant differences in mRNA expression levels between the two KMO variants (Figure 2A). Levels of t*KMO* mRNA expression were used as a control to normalise the expression levels of the KMO-Arg/Cys452 variants. As expected, *KMO* mRNA expression was not detected in non-transfected HEK293T cells or cells transfected with RFP as these cells do not express endogenous *KMO* (www.proteinatlas.org).

**Figure 2.**
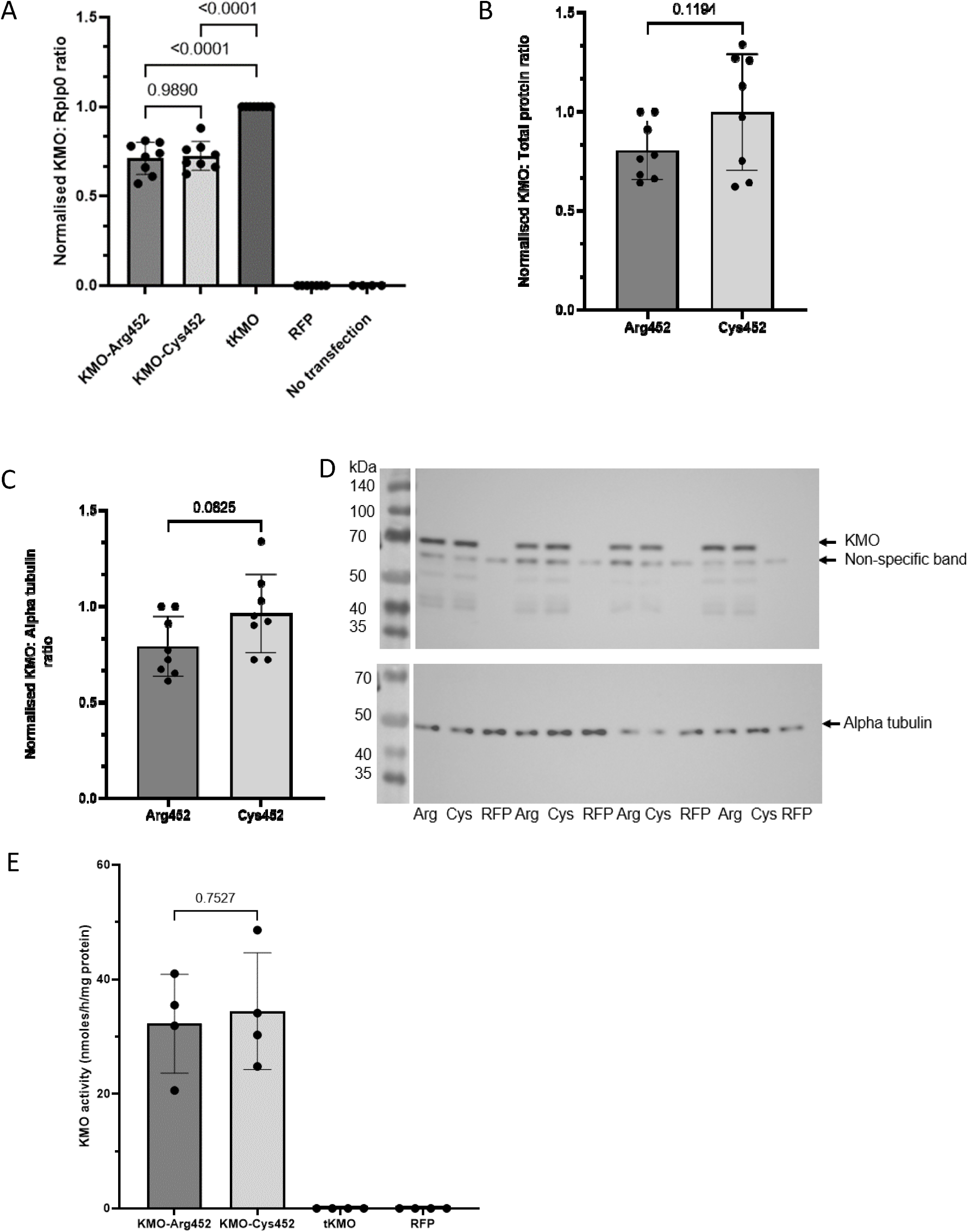
KMO-Arg452 and KMO-Cys452 mRNA and protein are expressed at similar levels in HEK293T cells and have similar activity levels. HEK293T cells were transfected to express KMO-Arg452-RFP, KMO-Cys452-RFP, tKMO-RFP or RFP alone. A) *KMO* mRNA expression was examined at 24 h post-transfection using RT-qPCR (n=8, One-Way ANOVA, Tukey’s test). KMO protein expression was examined at 48 h post-transfection using immunoblot analysis for KMO and normalised to B) total protein or C) alpha tubulin (n=8, unpaired t-test). KMO:total protein or KMO:alpha tubulin ratios were normalised to the first Arg452 sample on each gel. D) Examples of KMO and alpha-tubulin immunoblots. Molecular weight ladders were imaged under white light and are shown alongside the blots. E) KMO activity of the cells was measured 48 h post-transfection by measuring 3-HK levels using HPLC (n=4, unpaired t-test).

KMO protein expression was examined in HEK293T at 48 h post transfection by immunoblot analysis and revealed no significant differences between the two KMO variants when compared to levels of total protein (Figure 2B) or the expression of the house-keeping gene alpha-tubulin (Figure 2C). The molecular weight of KMO (50 kDa) in conjunction with RFP (25 kDa) was observed as a band of the expected size (∼70 kDa) (Figure 2D).

No significant differences in KMO enzymatic activity were observed between the KMO-Arg452 and KMO-Cys452 variants (Figure 2E). As expected, the truncated form of KMO (tKMO) had no enzymatic activity because it lacks the second putative transmembrane domain which is required for KMO activity [40].

### KMO-Arg452 protein exhibits reduced stability compared to KMO-Cys452

Inhibition of protein translation using cycloheximide (CHX) is a common approach to assess protein stability in cells [41]. HEK293T cells expressing the KMO-Arg452 or Cys452 variants were treated with CHX (50 µg/ml) or vehicle control (0.25 % (v/v) DMSO) for 4 or 8 h to examine the influence of the Arg/Cys452 substitution on KMO protein stability. Immunoblot analysis of cells treated with CHX revealed significant reductions in levels of KMO-Arg452 at 8 h compared to the respective untreated control at 0 h (0 h untreated vs. 8 h CHX *p* = 0.0108) (Figure 3). Reduced levels of KMO-Arg452 were also observed in cells treated with DMSO compared to the untreated control (Figure 3), possibly due to DMSO-induced cellular stress promoting the degradation of KMO [42]. In contrast, less pronounced, non-significant reductions in KMO-Cys452 levels were observed in cells treated with CHX or DMSO for 4 or 8 h compared to KMO-Cys452 levels in untreated cells at 0 h (Figure 3). Together these data indicate that the KMO-Arg452 variant has reduced protein stability or a faster turnover rate compared to KMO-Cys452.

**Figure 3.**
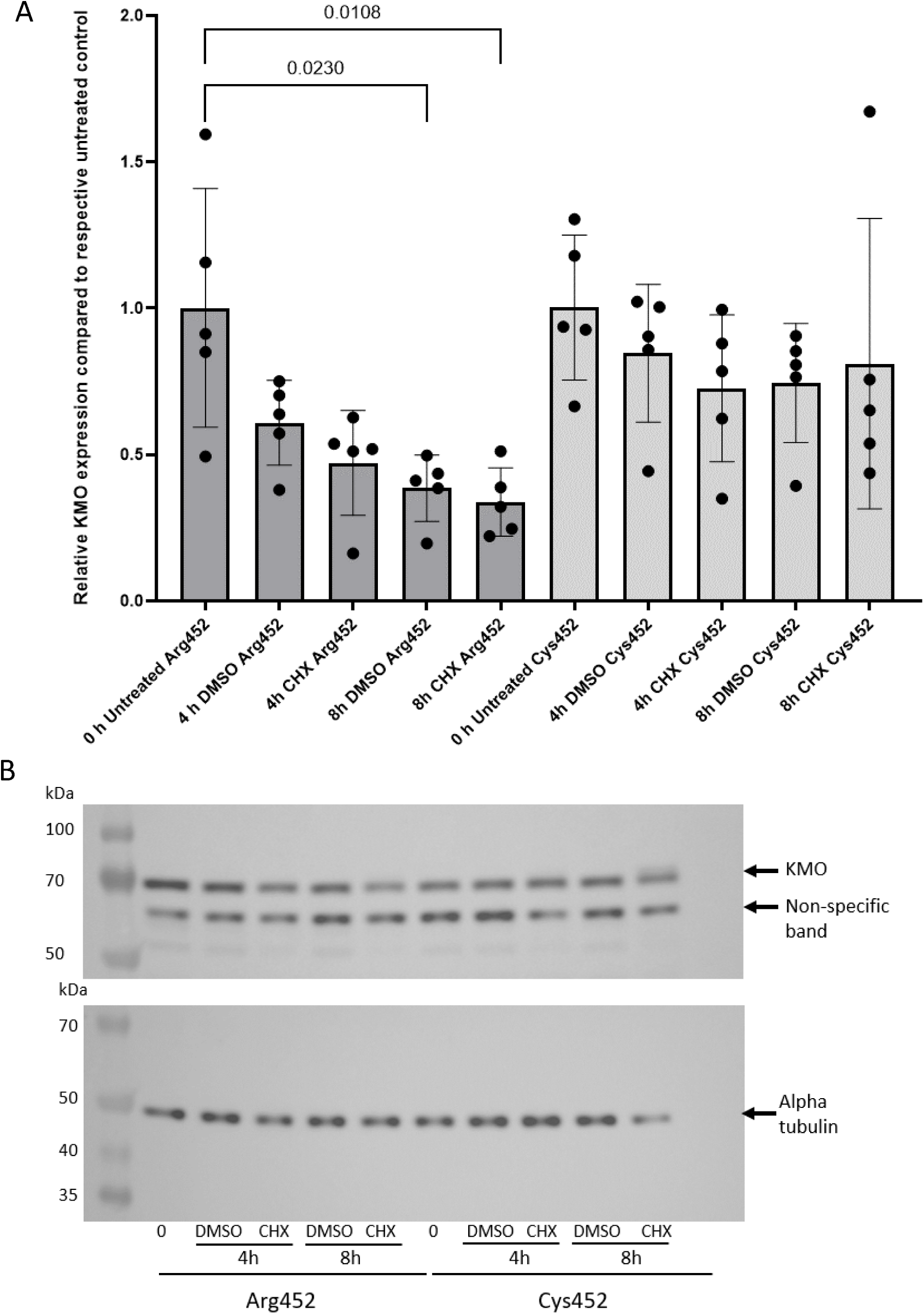
KMO-Arg452 protein stability is reduced compared to KMO-Cys452. HEK293T cells were transfected to express KMO-Arg452-RFP or KMO-Cys452-RFP. At 48 h post-transfection cells were treated with cycloheximide (CHX) (50 µg/ml) or 0.25 % (v/v) DMSO as vehicle control and incubated for 4-8 h. Cells were lysed and examined by immunoblot analysis for KMO and alpha tubulin. A) Results from densitometry analysis of KMO and alpha tubulin. KMO:alpha tubulin ratios for Arg452 and Cys452 samples were normalised to the average of the respective untreated control values for each KMO variant (n = 5 ± SD, One-Way ANOVA, Tukey’s multiple comparisons test). B) Examples of immunoblots for KMO and alpha tubulin.

### KMO-Arg452 and KMO-Cys452 localise to mitochondria to similar extents in HEK293T cells

Examination of cells using confocal microscopy revealed that both KMO-Arg452-RFP and KMO-Cys452-RFP co-localised with the mitochondrial marker HtrA2 in HEK293T cells, indicating that both variants localised to mitochondria (Figure 4A). The degree of colocalisation was confirmed by calculating Pearson’s and Mander’s coefficients to indicate overlap between KMO-RFP variants (red) and HtrA2 (green). Both KMO-Arg452 and KMO-Cys452 had similar Pearson’s coefficients (0.6 ± 0.09 for Arg452 and 0.58 ± 0.08 for Cys452) (Figure 4B), and Mander’s coefficients (0.36 ± 0.19 for Arg452 and 0.39 ± 0.19 for Cys452) (Figure 4C), and statistical analyses showed no significant differences between the two KMO variants for either coefficient. tKMO, which lacks a critical C-terminal domain necessary for localisation to the mitochondrial outer membrane [40], served as a negative control for KMO localisation to mitochondria as described previously [26]. As expected, tKMO did not localise to mitochondria but was distributed throughout the cytoplasm (Figure 4A). The Pearson’s and Mander’s coefficients for tKMO were lower than full length KMO (0.21 ± 0.08 and 0.11 ± 0.09 respectively), indicating a lack of co-localisation with HtrA2 (Figures 4B&C), and were significantly different compared to the KMO-Arg452 and KMO-Cys452 for both the Pearson’s coefficients (*p* = <0.0001 for tKMO vs. Arg452 and Cys452) and Mander’s coefficients (*p* = <0.0001 for tKMO vs. Arg452 and Cys452).

**Figure 4.**
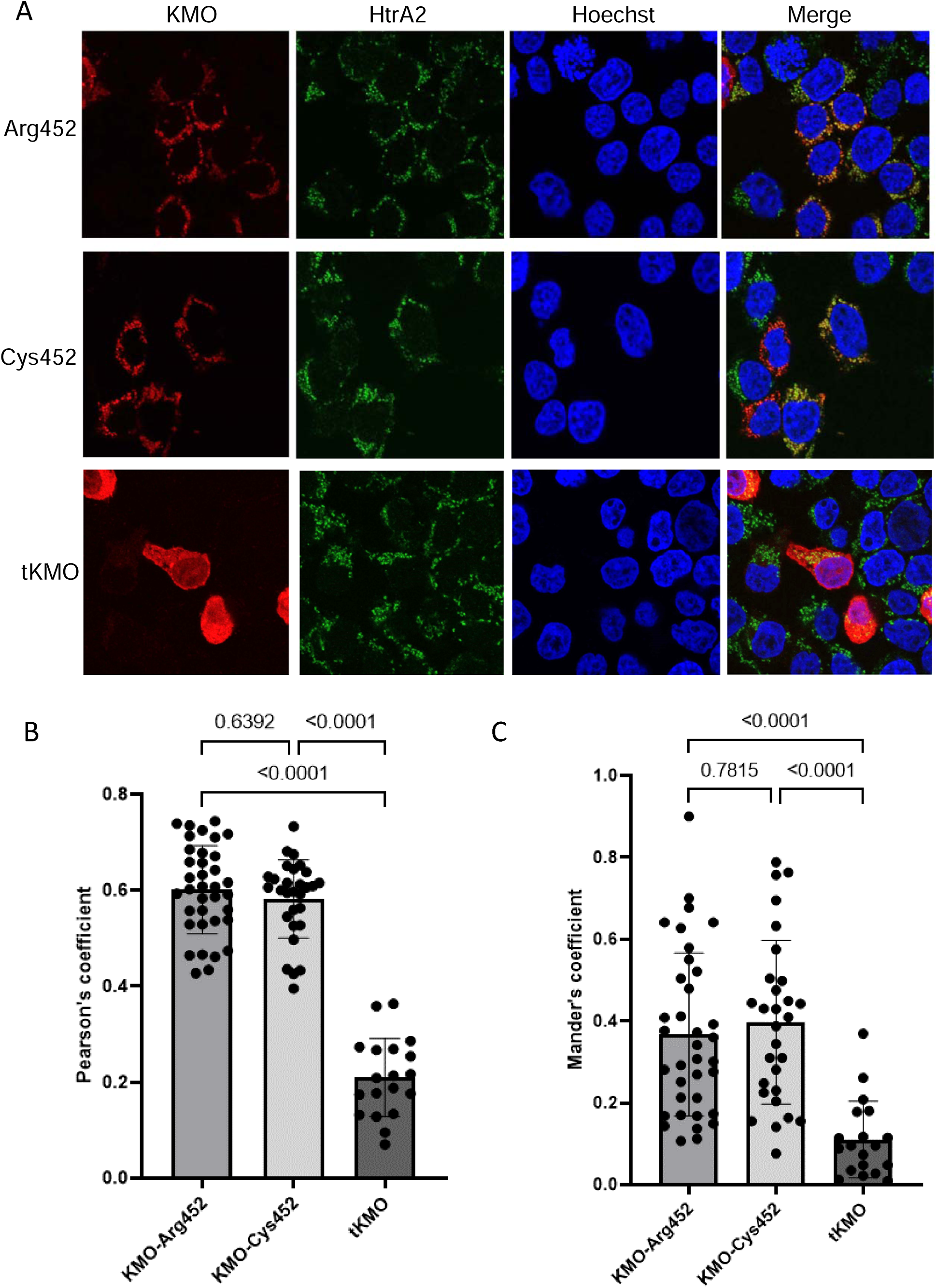
KMO-Arg452 and KMO-Cys452 both localise to mitochondria to similar extents. HEK293T cells were transfected to express KMO-Arg452-RFP, KMO-Cys452-RFP or tKMO-RFP. Cells were fixed and labelled with anti-HtrA2 as a mitochondrial marker and the nuclei stained using Hoechst. Cells were imaged using an Olympus FV1000 confocal microscope. A) KMO-Arg452-RFP, KMO-Cys452-RFP and tKMO-RFP co-localisation with HtrA2 in HEK293T cells. B) Pearson’s coefficient of the correlation between red and green pixels. C) Mander’s coefficient of colocalization of red and green pixels. Data are from three independent experiments and 19-36 cells per group were analysed. One-way ANOVA, Tukey’s post-hoc test.

### Molecular dynamics simulations predict reduced structural stability of KMO-Arg452 compared to the KMO-Cys452 variant

AlphaFold was used to assess structural differences between the KMO-Arg/Cys452 variants. The AlphaFold prediction data indicated that the C-terminal region of KMO is highly flexible and may adopt different conformations or structural arrangements (Figure S3 in Supplemental Data File 1). Such flexibility is likely to be associated with protein stability as well as interactions with potential protein partners. To gain further mechanistic insights, molecular dynamics (MD) simulations were performed. The MD simulation results showed that the root-mean-square deviation (RMSD) values of the protein complexes remained within approximately 2 Å during the later stages of the simulation (Figure 5A), indicating overall structural convergence. Notably, more pronounced RMSD fluctuations were observed for the Arg452 variant, suggesting reduced structural stability compared with the Cys452 variant. Consistent with the RMSD analysis, the Arg452 variant exhibited generally higher root-mean-square fluctuation (RMSF) values across the entire structure, with particularly pronounced fluctuations at the C-terminal region (Figure 5B). This result indicates increased overall flexibility and especially high mobility of the protein tail in the Arg452 variant. In addition, the Arg452 variant displayed a slightly larger radius of gyration (Rg) than the Cys452 variant (Figure 5C), indicating a more expanded and less compact structure. Consistently, the Arg452 variant exhibited higher solvent-accessible surface area (SASA) values throughout the simulation (Figure 5D), particularly at later stages, reflecting a more exposed structural organization. Structural comparison of the initial and final conformations further supports this conclusion. The cyan structures in Figure 5E represent the initial conformations of the KMO variants, while the light magenta structures correspond to the stable conformations obtained at the end of the 300 ns simulation (Figure 5E). In the Cys452 variant, the C-terminal region shifted toward the protein core during the simulation, resulting in reduced tail flexibility at the later stages. Collectively, these MD simulation results suggest that the Arg452 variant is more flexible and less structurally stable than the Cys452 variant.

**Figure 5.**
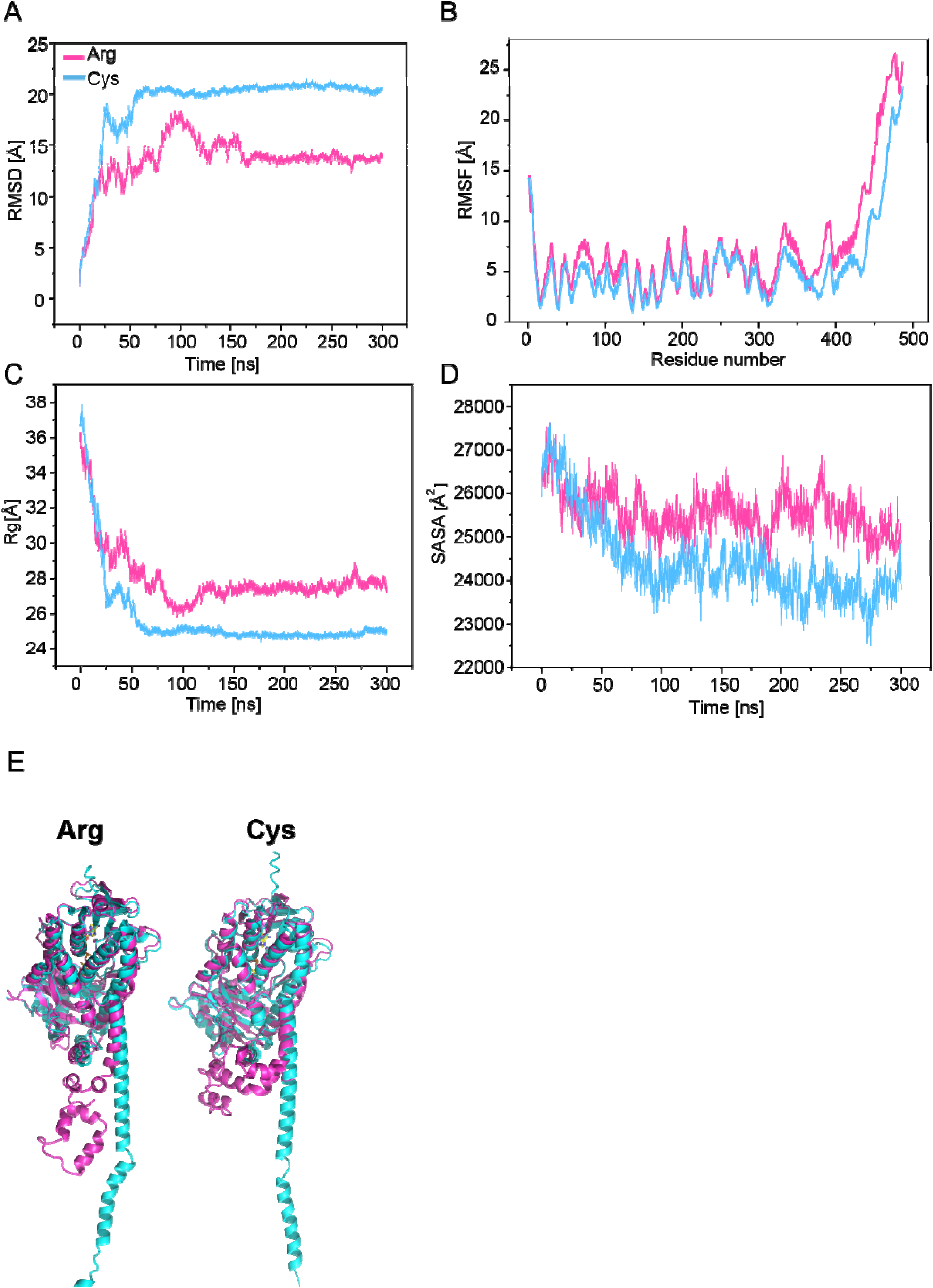
KMO variant molecular dynamics (MD) simulation results. (A) Root mean square deviation (RMSD) analysis of protein complex. (B) Root mean square fluctuation (RMSF) analysis of protein complex. (C) The radius of gyration (Rg) analysis of protein complex. (D) Solvent accessible surface area (SASA) values of protein complex. E) Initial (cyan) and stable (light magenta) conformations after 300 ns of simulation for KMO variants, showing a C-terminal shift toward the protein core and reduced tail flexibility in the Cys452 variant.

### Immunoprecipitation of KMO-Arg452 and KMO-Cys452 reveals differential KMO-interacting proteins

Since the AlphaFold and MD simulation data indicated differences in the flexibility and structural stability of the C-terminus between the KMO variants, we examined whether this could alter protein-protein interactions. Immunoblot analysis of immunoprecipitated samples confirmed that both KMO-Arg452 and KMO-Cys452 were pulled-down from cell lysates using RFP-trap beads (Figure 6A). Mass spectrometry was then used to analyse proteins that co-immunoprecipitated with KMO-Arg452-RFP or KMO-Cys452-RFP in three independent sets of cells. In agreement with the immunoblot analyses in Figures 2B&C, the normalised abundance data from the mass spectrometry analysis showed that the expression levels of KMO-Arg452 and KMO-Cys452 were similar (KMO-Arg452 normalised abundance = 31.2 ± 19.1, KMO-Cys452 normalised abundance = 27.7 ± 20.9, n=3). Proteins that specifically interacted with the KMO variants and had a confidence score ≥10 were then examined using Venn diagram analysis. Strikingly, this analysis revealed 328 proteins that specifically interacted with the KMO-Arg452 variant, 555 proteins that specifically interacted with KMO-Cys452, and only 156 proteins that interacted with both KMO variants (Figure 6B). To increase the stringency of the data, only proteins with confidence scores ≥40 that specifically interacted with the Arg452 or Cys452 KMO variants were analysed further (see Table S1 in Supplemental Data File 1). Peptide counts, confidence values and normalised abundance data for all proteins identified by mass spectrometry that interacted with KMO-Arg452 or KMO-Cys452 (with RFP-interacting proteins removed) are shown in Supplemental Data File 2.

**Figure 6.**
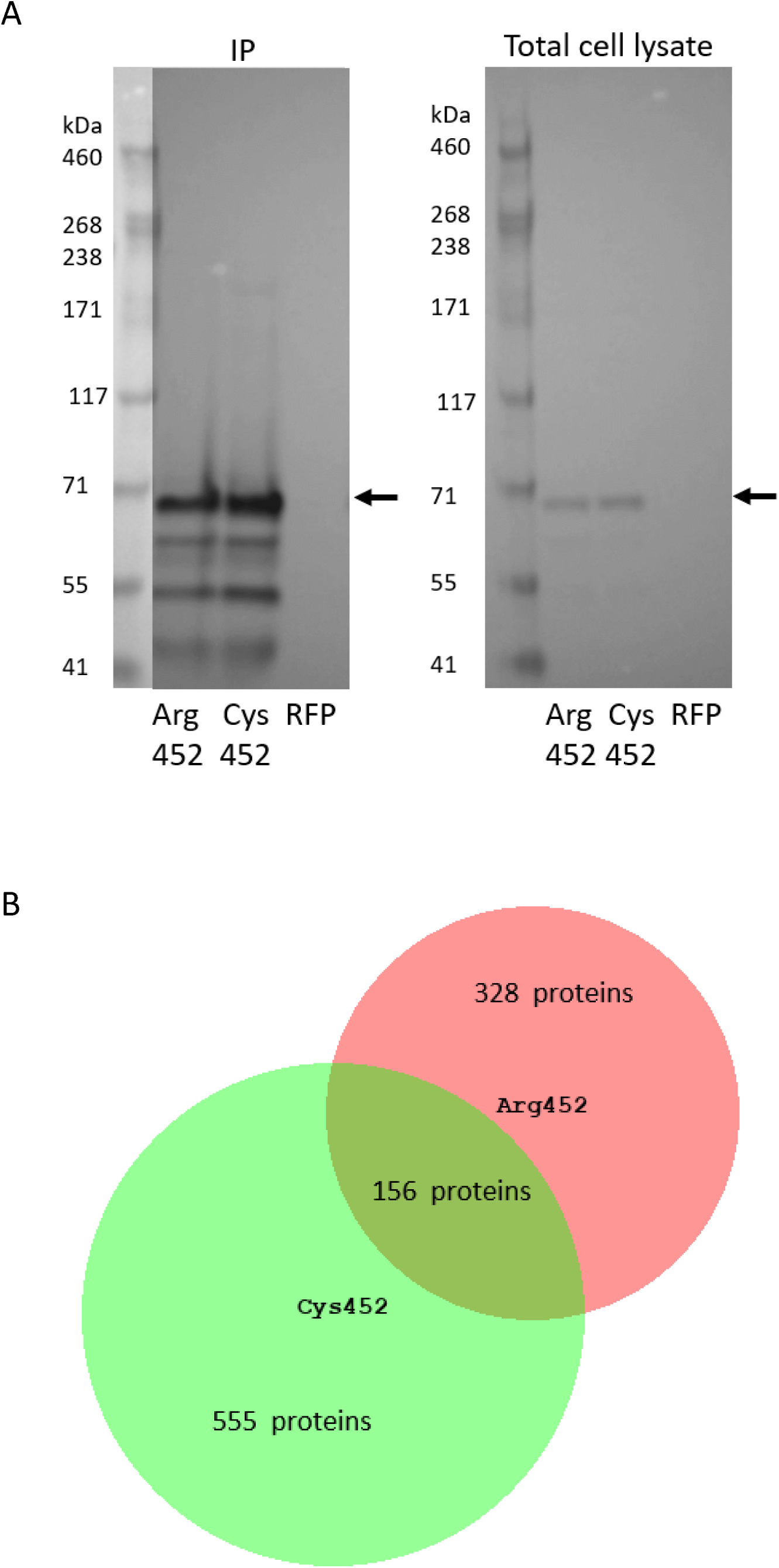
RFP-trap pull-downs of KMO variants and interacting proteins. HEK293T cells were transfected to express KMO-Arg452-RFP, KMO-Cys452-RFP or RFP alone. A) KMO-RFP variants were pulled-down from cell lysates using RFP-trap agarose beads (IP) and examined alongside total cell lysates by immunoblot analysis using an anti-KMO antibody. Arrows indicate KMO-RFP. B) Mass spectrometry was used to examine KMO-Arg/Cys452-interacting proteins in three separate sets of cells with each biological replicate examined by mass spectrometry in three technical repeats. Interacting proteins with confidence scores >10 were analysed using the Bio Venn software and revealed proteins that specifically interacted with the two KMO variants and proteins that interacted with both KMO variants.

### GO-term enrichment analysis and STRING protein interaction network analysis of KMO-Arg452 interacting proteins reveals enrichment of terms related to microtubules and muscle structure

GO-term enrichment analysis of KMO-interacting proteins with confidence scores ≥40 revealed proteins that interacted with KMO-Arg452 had significant enrichment of cellular component terms associated with microtubules such as cortical microtubule cytoskeleton, microtubule end, and kinesin complex (Figure 7A and Table S2 in Supplemental Data File 1). The top ten cellular component terms for KMO-Arg452-interacting proteins also showed enrichment of terms relating to muscle structure such as A band, myosin complex, contractile muscle fibre, sarcomere and myofibril (Figure 7A and Table S2 in Supplemental Data File 1). Lists of KMO-Arg452-interacting proteins associated with selected GO terms of interest are shown in Table S3 in Supplemental Data File 1. No significant biological process terms were observed for KMO-Arg452 interacting proteins. STRING protein-protein interaction network analysis was used to further examine associated functions of groups of proteins that interacted with KMO-Arg452. Although the protein-protein interaction enrichment *p*-value was not significant (*p* = 0.089) due to the small number of protein interactors, this analysis revealed clusters of proteins involved in microfilament motor activity, kinetochore binding and ribosomal proteins (Figure 7B).

**Figure 7.**
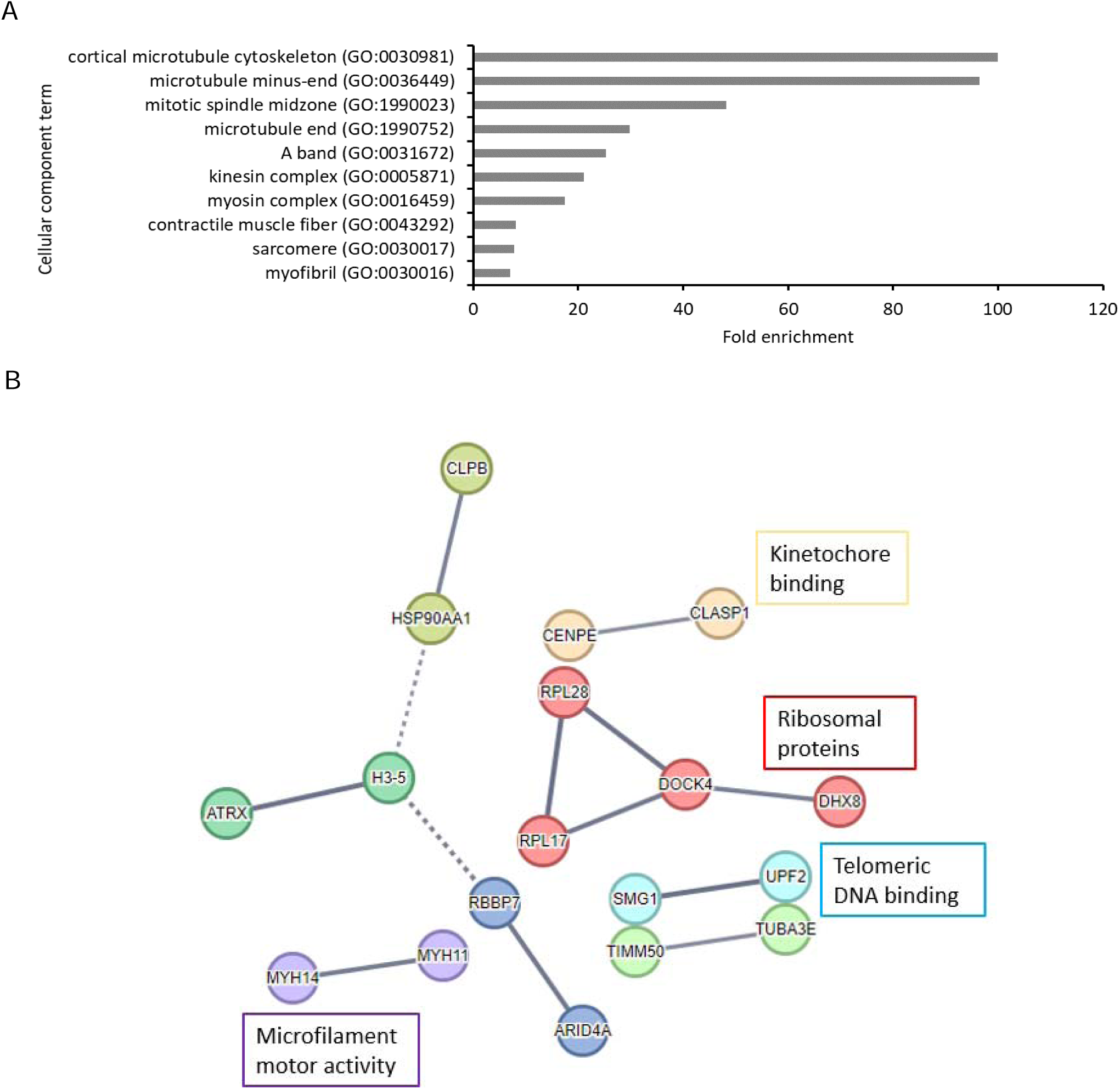
GO-term enrichment analysis and STRING analysis of KMO-Arg452 interacting proteins. KMO interacting proteins specific to the Arg452 variant and with confidence scores >40 were analysed using A) GO-term enrichment analysis to derive cellular component terms associated with KMO-Arg452 interacting proteins (top ten GO terms are shown) and B) STRING protein interaction network analysis to derive networks associated with KMO-Arg452 interacting proteins. Network edges show confidence where the thickness of the lines represent the strength of the supporting data.

### GO-term enrichment analysis and STRING protein interaction network analysis of KMO-Cys452 interacting proteins reveals enrichment of terms for synaptic function

Similarly to the KMO-Arg452 variant, KMO-Cys452 interacting proteins were enriched in cellular component terms related to microtubules but also included actin related terms such as contractile actin filament bundle and stress fibre (Figure 8A and Table S4 in Supplemental Data File 1). In contrast to the KMO-Arg452 variant, however, proteins that interacted with KMO-Cys452 showed a high enrichment of proteins associated with synapse structure and synaptic function. For example, the top ten cellular component terms included a 16.6-fold enrichment of proteins associated with “terminal bouton” (Figure 8A and Table S4 in Supplemental Data File 1), whereas analysis of the top ten biological process terms revealed a 29.3-fold enrichment of proteins associated with the biological process term “regulation of synaptic vesicle recycling”, and a 5.6-fold enrichment of proteins involved in synaptic organisation (Figure 8B and Table S4 in Supplemental Data File 1). Other biological process terms enriched for KMO-Cys452 interacting protein included the regulation of autophagy, regulation of actin filament-based process, and regulation of cellular component organisation (Figure 8B and Table S4 in Supplemental Data File 1). Lists KMO-Cys452-interacting proteins associated with selected GO terms of interest are shown in Table S5 in Supplemental Data File 1. STRING analysis of proteins that specifically interacted with KMO-Cys452 revealed an interaction network with a significant enrichment in protein-protein interactions (*p* = 0.00085) and 29 edges compared to the expected 15 edges, indicating that enrichment of proteins into biologically associated clusters was not a random event. This included enrichment of protein clusters involved in microtubule plus-end binding, actin cytoskeleton, centriole-centriole cohesion, and phosphatidylinositol signalling (Figure 8C).

**Figure 8.**
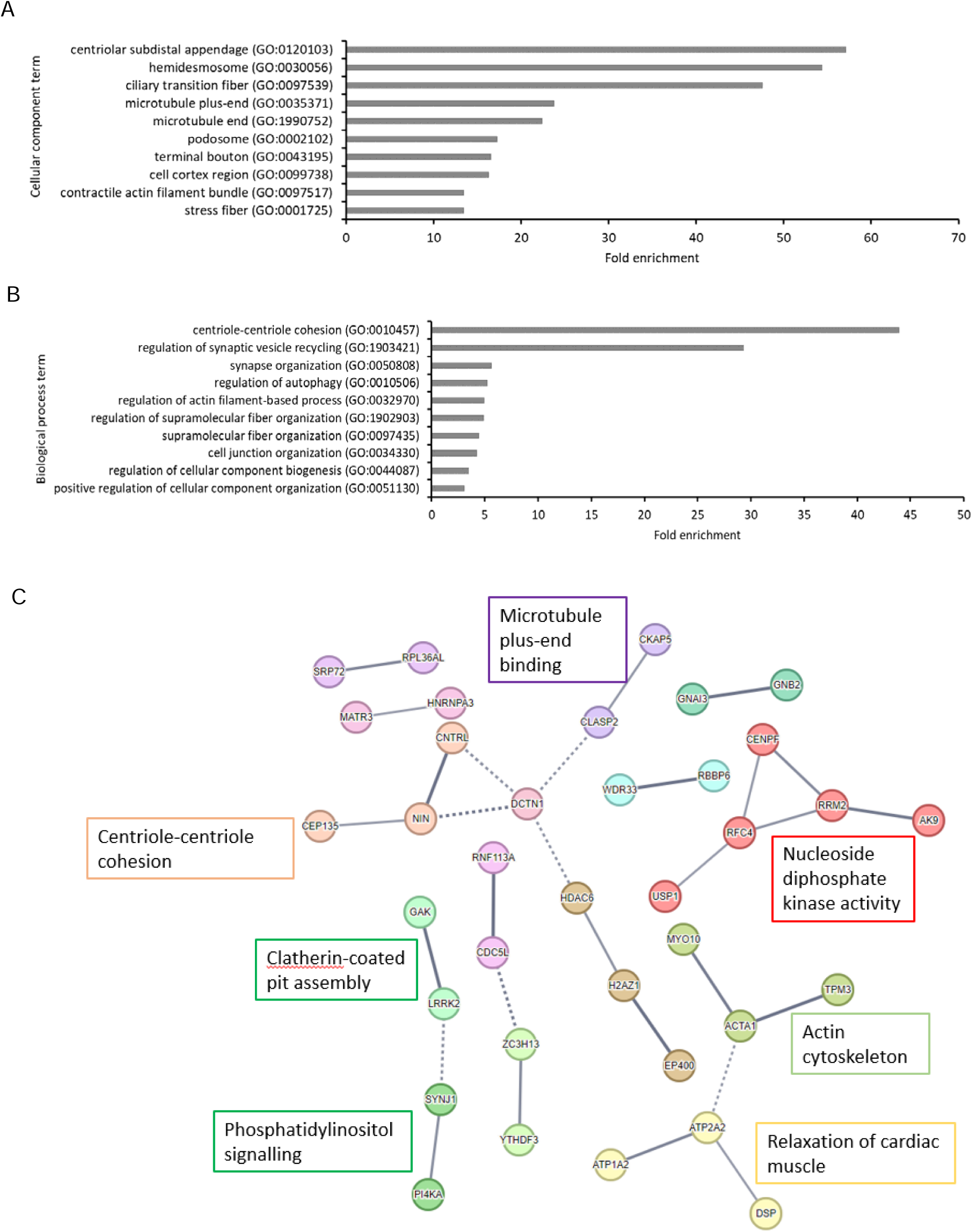
GO-term enrichment analysis and STRING analysis of KMO-Cys452 interacting proteins. KMO interacting proteins specific to the Cys452 variant and with confidence scores >40 were analysed using A) GO-term enrichment analysis to derive cellular component terms associated with KMO-Cys452 interacting proteins (top ten GO terms are shown), B) biological process terms associated with KMO-Cys452 interacting proteins (top ten GO terms are shown), and C) STRING protein interaction network analysis to derive networks associated with KMO-Cys452 interacting proteins. Network edges show confidence where the thickness of the lines represent the strength of the supporting data.

### GO-term enrichment analysis and STRING protein interaction network analysis of shared KMO-Arg452/Cys452 interacting proteins reveals enrichment of terms related to muscle function and the cytoskeleton

A list of proteins that interacted with both KMO-Arg452 and KMO-Cys452 and had confidence values ≥40 is shown in Table S6 in Supplemental Data File 1. GO analysis of these proteins showed enrichment of cellular component terms associated with muscle structure (Figure S4A and Table S7 in Supplemental Data File 1), and biological process terms related to muscle function, cytoskeletal organisation and organelle organisation (Figure S4B and Table S7 in Supplemental Data File 1). STRING protein interaction network analysis of these shared protein interactors revealed a network with an enrichment of protein-protein interactions (*p* = 0.0029) and 13 edges compared to the expected 5 edges, indicating enrichment of proteins into biologically associated clusters was not due to chance. This included enrichment of protein clusters associated with microtubules, sarcoplasmic reticulum membrane calcium release and protein localisation to M band (Figure S4C in Supplemental Data File 1).

### KMO rs1053230 genotype is associated with altered clinical features in a schizophrenia cohort

Finally, we sought to determine whether the KMO rs1053230 genotype was associated with key clinical features relevant to schizophrenia in a cohort of 156 individuals with schizophrenia and 170 healthy controls via a set of clinical assessments and cognitive tests for processing speed and working memory. Characteristics of the cohort are shown in Table S8 in Supplemental Data File 1. The genotypes for rs1053230 were determined in the cohort and as expected found to be in Hardy-Weinberg equilibrium (Χ^2^ = 0.02, *p* = 0.89). The Digit Symbol Coding task assesses processing speed, a core cognitive modality related to mental efficiency with simple tasks and indicated a significant effect of rs1053230 genotype across the entire cohort (F(1,307) = 4.25, *p* = 0.040), but no genotype × diagnosis effect was observed (F = 0.05, *p* = 0.82). Carriers of the minor allele encoding KMO-Cys452 had significantly greater performance on the processing speed task (M = 68.4 ± 19.0) compared to homozygotes for the major allele (M = 67.1 ± 20.3). No significant effects of genotype on working memory were observed (F = 0.16, *p* = 0.69) or genotype × diagnosis (F=0.12, *p* = 0.73) in the cohort. Assessment of longitudinal or trait depression via the MTSD found a significant effect of genotype, with carriers of the minor allele encoding KMO-Cys452 reporting greater symptoms of longitudinally experienced depression (M = 19.2 ± 16.6) than major allele homozygotes (M = 14.2 ± 14.4; F(1,312) = 4.28, *p* = 0.039). However, no genotype × diagnosis interaction was observed for this trait. For recent or state depression, a non-significant but similar trend was found for more recent depressive symptoms in the minor allele carrier group (F=2.97, *p* = 0.09), again without diagnosis × genotype interaction. Among patients, there was no effect of genotype on total psychiatric symptoms as measured with the BPRS (F = 0.75, *p* = 0.39) or negative symptoms as measured with BNSS (F = 1.79, *p* = 0.18). In summary, rs1053230 genotype is not associated with positive or negative symptoms of psychosis but is associated with chronic, trait-like depressive mood symptoms in schizophrenia patients.

## Discussion

This study examined the KMO SNPs rs2275163 and rs1053230 using cell-based *in vitro* models to further understand the underlying mechanisms of how these SNPs may affect KMO expression and function. Initially we examined the effects of rs2275163 genotype on *KMO* mRNA expression, since this SNP has been previously associated with lower predictive pursuit and more visuospatial errors in individuals with the C/C genotype compared to the C/T genotype [12]. Wonodi *et al*. (2011) [12] also found that *KMO* mRNA expression was slightly lower in the frontal cortex of schizophrenia patients with the rs2275163 C/C genotype compared to the C/T genotype, although this was not a statistically significant difference. Our data showed that the rs2275163 genotype had no observable effect on *KMO* pre-mRNA stability or alternative splicing of *KMO* mRNA. This is in agreement with a study by Johansson *et al*. (2013) [43] which did not observe any significant effect of rs2275163 genotype on *KMO* mRNA expression in fibroblasts from patients with schizophrenia, suggesting that this SNP does not directly affect *KMO* expression. rs2275163 genotype has also not been associated with a diagnosis of schizophrenia [12,44]. The effect of rs2275163 genotype on schizophrenia endophenotypes [12] may therefore be due to other factors such as linkage disequilibrium of rs2275163 with other causative genetic variants.

We next examined the influence of the genotype of the coding SNP rs1053230 on KMO expression and function. This followed previous observations that altered KMO expression and activity in patients with schizophrenia and bipolar disorder were associated with the genotype of this SNP [22,45]. In particular, the KMO-Arg452 variant has been associated with reduced *KMO* expression in both lymphoblastoid cells and in the hippocampus of epileptic patients [22]. KMO-Arg452 has also been associated with increased levels of KYNA in the CSF of patients with bipolar disorder [22]. Finally, in a combined group of controls and schizophrenia patients, Arg452 homozygotes (CC) were shown to have reduced cognitive function compared to individuals with the CT or TT genotypes [23], suggesting that reduced KMO expression due to the rs1053230 C genotype may lead to increased KYNA and cognitive deficits. Here we partially replicated lower cognitive performance associated with homozygous C genotype. However, we did not observe any significant differences in mRNA expression or protein expression in HEK293T cells expressing the KMO-Arg/Cys452 variants. In agreement with our study, previous reports have found no effect of rs1053230 genotype on *KMO* mRNA expression in fibroblasts from schizophrenia patients [43] or in the frontal eye field of patients [12].

The enzymatic activity of KMO requires the presence of the second transmembrane domain located at the C-terminus of KMO [40]. Since the rs1053230 genotype has been associated with levels of KYNA in the CSF [22,45], we examined whether the KMO-Arg/Cys452 substitution would affect KMO enzymatic activity. However, we did not observe any significant differences in 3-HK production in cells expressing the KMO-Arg/Cys452 variants, suggesting that rs1053230 genotype does not affect the enzymatic activity of KMO in HEK293T cells. This agrees with a study by Wonodi *et al.* (2011) [12] which found no significant association of rs1053230 genotype with KMO activity in the brain of schizophrenia patients. The targeting of KMO to the outer mitochondrial membrane has been shown to be dependent on two arginine residues (Arg461 and Arg462 in pig KMO) within the C-terminus of KMO in close proximity to Arg/Cys452 [40]. However, we demonstrated here that both the KMO-Arg452 and KMO-Cys452 variants localised to mitochondria to a similar extent, indicating that this SNP does not affect the cellular localisation KMO in HEK293T cells.

Interestingly, inhibition of protein synthesis using cycloheximide revealed that the KMO-Arg452 variant had reduced protein stability compared to KMO-Cys452, indicating that protein turnover rates of the two variants, rather than protein expression, are affected by the rs1053230 genotype. Arginine is a hydrophilic, positively charged amino acid, whereas cysteine is hydrophobic. The different properties of these amino acids may influence protein structure and therefore the stability of KMO. In support of this hypothesis, AlphaFold modelling and MD simulations predicted that the KMO-Arg452 variant has increased flexibility and reduced stability of the C-terminus compared to the Cys452 variant.

Alternatively, protein turnover differences between the two KMO variants may be due to post-translational modifications in the C-terminus of KMO influenced by the rs1053230 genotype. For example, arginine residues in the RXR motif can be methylated [46] leading to downstream post-translational modifications such as ubiquitination [47]. Therefore, it is possible that residue Arg452 is methylated leading to KMO degradation through the ubiquitin proteasome pathway, although this requires further investigation.

Since AlphaFold and MD simulations predicted differences in the structure and flexibility of the KMO C-terminus between the two variants, we next examined whether KMO-Arg/Cys452 substitution influences KMO-protein interactions. Interestingly, we observed differences in KMO-interacting proteins between the two variants and identified 328 proteins that specifically interacted with KMO-Arg452, and 555 that interacted with KMO-Cys452. We confirmed our previous findings that KMO interacts with the large scaffolding protein huntingtin (HTT) [26], although the confidence score was <40 and was therefore not included in the GO analysis. We also observed pull-down of Disrupted in Schizophrenia-1 (DISC1) (confidence score 22.6) which has previously been shown to interact with KMO [48], further confirming successful co-purification of KMO and protein interactors. The interaction of KMO with DISC1 is also of interest because of the association of this protein with schizophrenia [49] and the role of DISC1 in the microtubule-mediated transport of mitochondria in neurons [50,51]. Due to the RFP-trap pull-down assay design it is possible that not all proteins pulled-down with KMO are direct interactors but may be part of protein complexes associated with KMO. However, we did not observe any significant enrichment of outer mitochondrial membrane proteins, indicating successful extraction of KMO from the outer mitochondrial membrane and specific pull-down of KMO and interacting proteins.

GO-term enrichment analysis revealed that both KMO variants had enrichment of cellular component terms related to microtubule function such as cortical microtubule cytoskeleton, microtubule minus-end, and kinesin complex for KMO-Arg452, and microtubule plus-end for KMO-Cys452. The microtubule system has roles in many cellular processes including vesicle and mitochondrial transport, cell division and cell migration. Since KMO is an outer mitochondrial membrane protein, it is possible that KMO-protein interactions have roles in mitochondrial transport on the microtubule system. We have previously shown that KMO has a role in mitochondrial dynamics which is independent of enzymatic activity [52]. Since there is an overlap between proteins involved in mitochondrial dynamics and mitochondrial transport [53,54] and KMO-interactors included components of the machinery used for mitochondrial transport such as tubulins, dynactin, and myosin, it would be interesting to examine whether interactions between KMO and the microtubule system have a role in mitochondrial transport.

Interestingly, KMO-Cys452-interacting proteins were highly enriched in GO terms linked to synaptic organisation and function such as regulation of synaptic vesicle recycling, terminal bouton and synaptic organisation, indicating that KMO-Arg/Cys452 variants may have functional differences in neurons. Although this study examined KMO protein interactors in HEK293T cells, neurons are known to express endogenous KMO [21]. KMO-Cys452-specific interactions with proteins involved in synaptic function and organisation suggest that KMO may have a role in neuronal development and/or function, with the Cys452 variant being more efficient in the relevant functional interactions. However, further studies would be required to elucidate the underlying mechanisms and whether KMO genotype can influence neurogenesis and contribute to the risk of developing schizophrenia along with other genetic and environmental factors. In support of these findings, Lavebratt *et al*. (2014) [22] found that patients with bipolar type I with psychosis during mania were less likely to have the KMO-Cys452 variant than the Arg452 variant compared to patients without psychosis or healthy controls. In contrast, our clinical data found no difference in total symptoms in a schizophrenia sample between the genotypes, but an increased experience of depressive symptoms in carriers of the Cys452 variant.

A potential limitation of this study is that the model used is not necessarily representative of *in vivo* situations, since HEK293T cells do not express endogenous KMO. It would therefore be important to examine the expression and function of these KMO variants in cells that express endogenous KMO such as microglia [20] and neurons [21]. Furthermore, this is a highly simplified model compared to the human brain where many genetic and environment factors may regulate levels of KMO expression and activity, particularly in patients with schizophrenia and bipolar disorder where there may be increased levels of oxidative stress [55,56,57] or inflammatory mediators [58, 59] which may also influence KMO expression and function.

In conclusion, we report the novel finding that KMO-Arg452 has reduced protein stability compared to KMO-Cys452. Notably, we find that the rs1053230 variants have differential protein-protein interactions that point to disparate involvement of the KMO-Arg/Cys452 variants in synaptic structure and function, the effects of which could have significant implications for rs1053230 genotype in neuronal development and associated psychiatric conditions.

## Acknowledgements

This work was supported by funding from the National Institute of Mental Health (Silvio O. Conte Center for Translational Mental Health Research) grant number MH-103222.

We thank the Advanced Imaging Facility (RRID:SCR_020967) at the University of Leicester for support and Dr Kees Straatman for assistance with confocal microscopy.

We also thank the Genomics Core facility at the University of Leicester for help and advice.

The mass spectrometry work was supported by the John and Lucille van Geest Foundation, and the National Institute for Health and Care Research Leicester Biomedical Research Centre. T.H.C. was funded by the National Institute for Health and Care Research Leicester Biomedical Research Centre.

This manuscript has been uploaded onto the bioRxiv pre-print server.

## Author contributions CRediT statement

Mary Collier: Investigation, Methodology, Formal Analysis, Visualisation, Project Administration, Conceptualization, Writing – original draft.

Georgia Ceeney: Investigation, Formal Analysis, Data curation, Writing – review and editing

Joshua Chiappelli: Investigation, Formal Analysis, Data curation, Resources, Project Administration, Writing – original draft (clinical data)

Sathyasaikumar Korrapati: Investigation, Formal Analysis, Writing – review and editing Thong Huy Cao: Data curation, Formal Analysis, Writing – review and editing

Paulene Quinn: Investigation, Writing – review and editing

Junfeng Ma: Investigation, Formal Analysis, Visualisation, Writing – original draft (molecular dynamics section)

Aashti Shauriq: Investigation

Nicolas Sylvius: Methodology, Formal Analysis, Writing – review and editing Edward J Hollox: Methodology, Writing – review and editing

Donald J L Jones: Supervision

Andrew Hudson: Supervision, Writing – review and editing

Elliot L Hong: Funding Acquisition, Conceptualization, Resources, Project Administration, Supervision, Writing – review and editing

Nigel Scrutton: Supervision, Writing – review and editing

Robert Schwarcz: Funding Acquisition, Conceptualization, Project Administration, Supervision, Writing – review and editing

Flaviano Giorgini: Funding Acquisition, Conceptualization, Project Administration, Supervision, Writing – review and editing

Data availability: All data is available on reasonable request.

