## Supplemental File 1 for "The single nucleotide polymorphism rs1053230 modulates kynurenine 3-monooxygenase stability and is associated with cognitive and mood phenotypes"

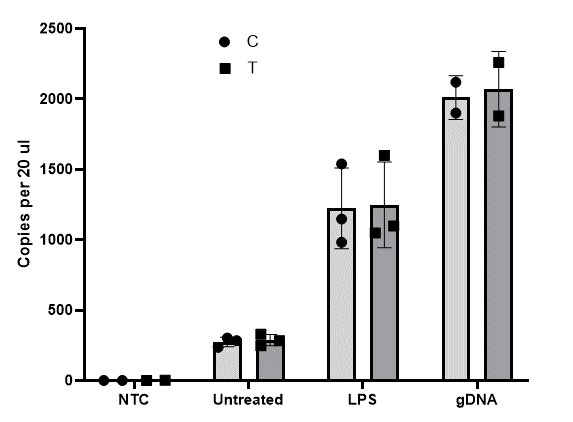

**Figure S1. rs2275163 genotype had no effect on KMO pre-mRNA stability.** THP-1 cells were treated with LPS (0.1 µg/ml) for 24 h to induce KMO expression or left untreated. RNA was isolated and the effect of rs2275163 genotype on KMO pre-mRNA stability was examined using ddPCR with rs2275163 C/T-specific MGB probes (n=3). NTC = no template control; gDNA = THP-1 genomic DNA.

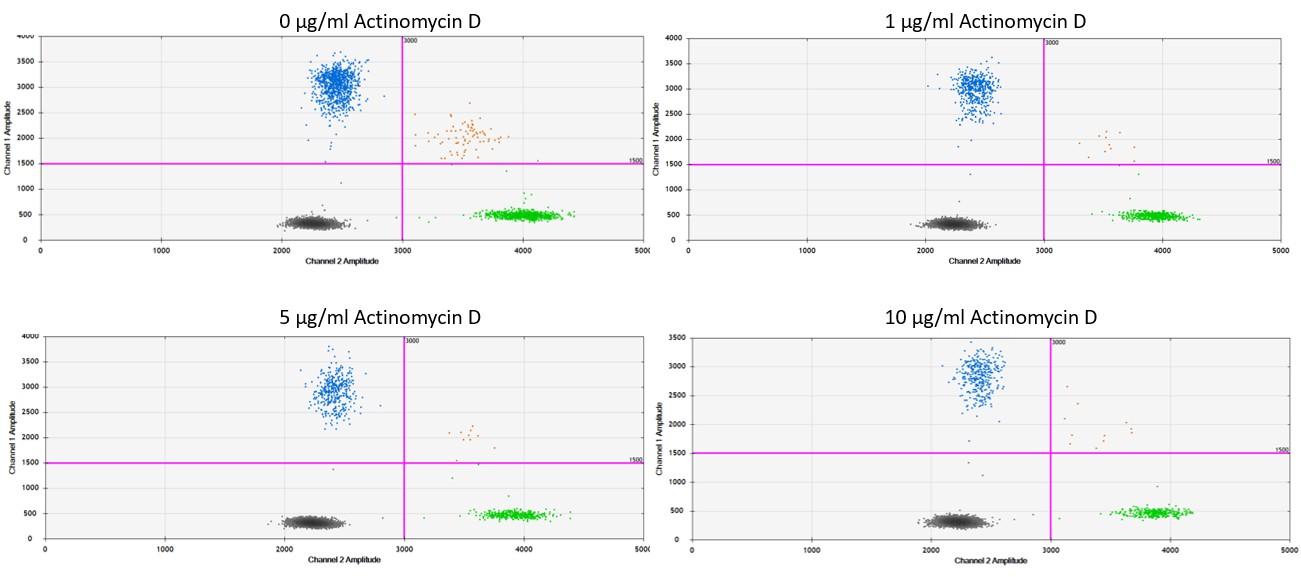

A

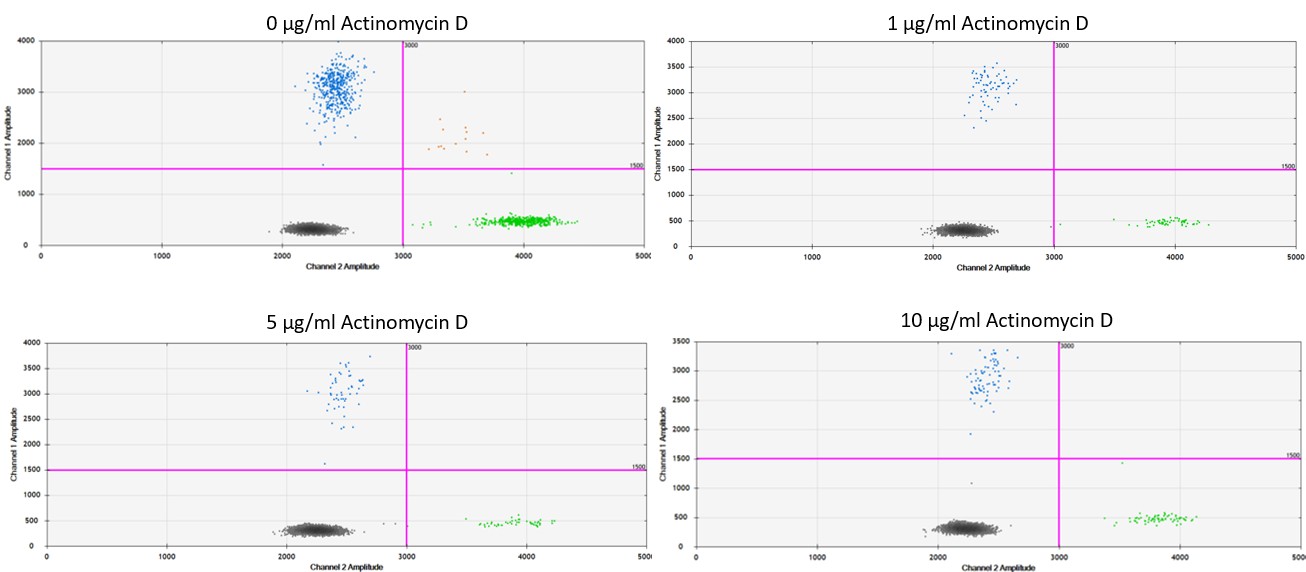

B

**Figure S2.** ddPCR dot plots of KMO pre-mRNA expression in THP-1 cells. THP-1 cells were treated with LPS (0.1 µg/ml) for 24 h to induce KMO expression. Cells were then treated with 1-10 µg/ml actinomycin D for 1 h (A) or 4 h (B). ddPCR with rs2275163 C/T-specific MGB probes was carried out on cDNA for each sample. Blue dots = C-FAM positive droplets; Green dots = T-HEX positive droplets; Orange dots = C+T positive droplets; Black dots = negative (empty) droplets.

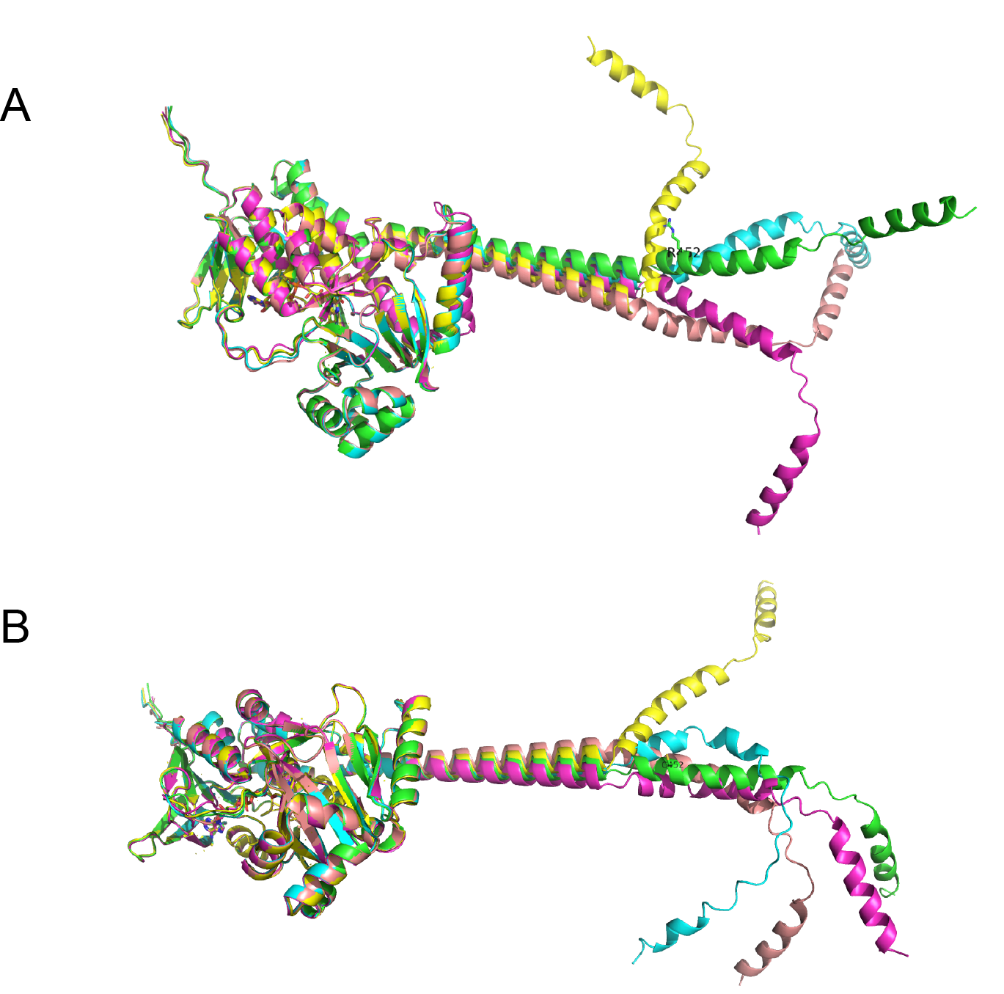

**Figure S3.** AlphaFold KMO variant structure prediction results. (A) The top 5 structures predicted from AlphaFold3 for variant Arg452. (B) The top 5 structures predicted from AlphaFold3 for variant Cys452.

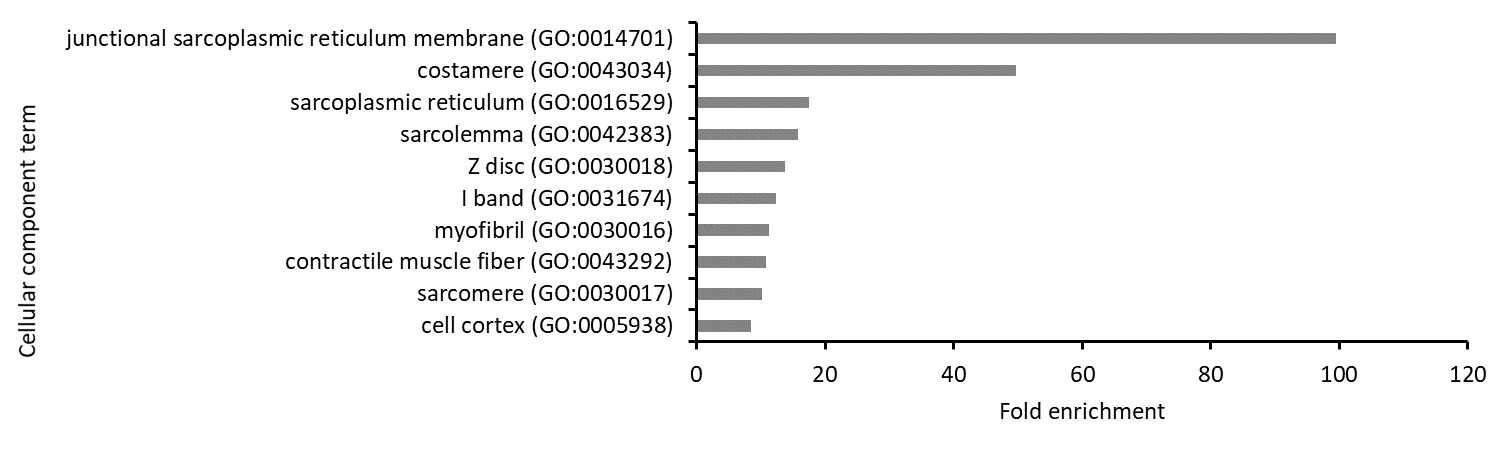

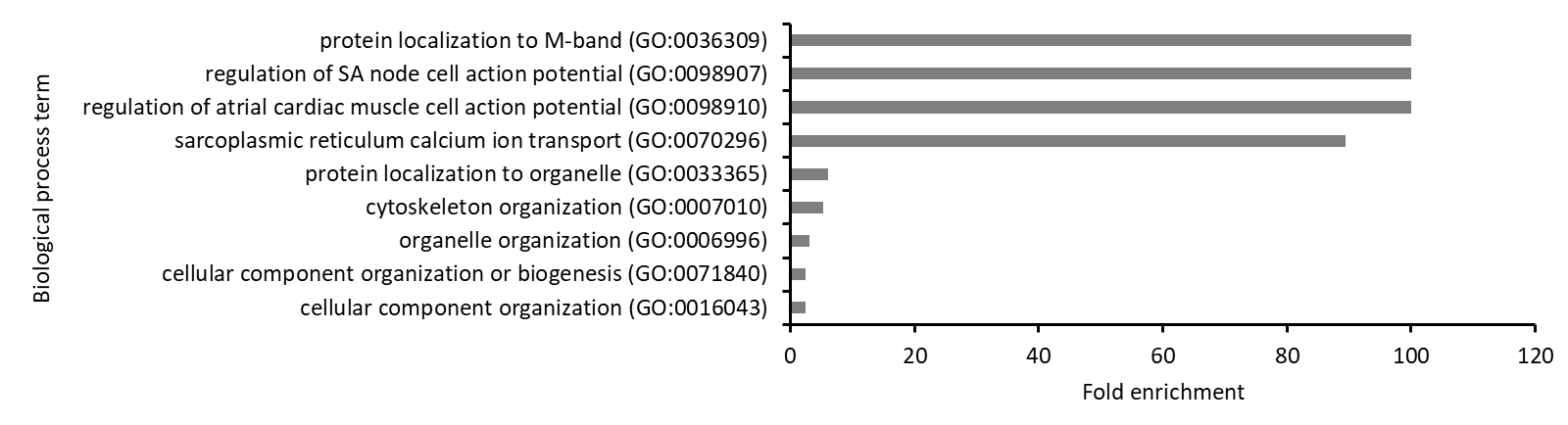

A

B

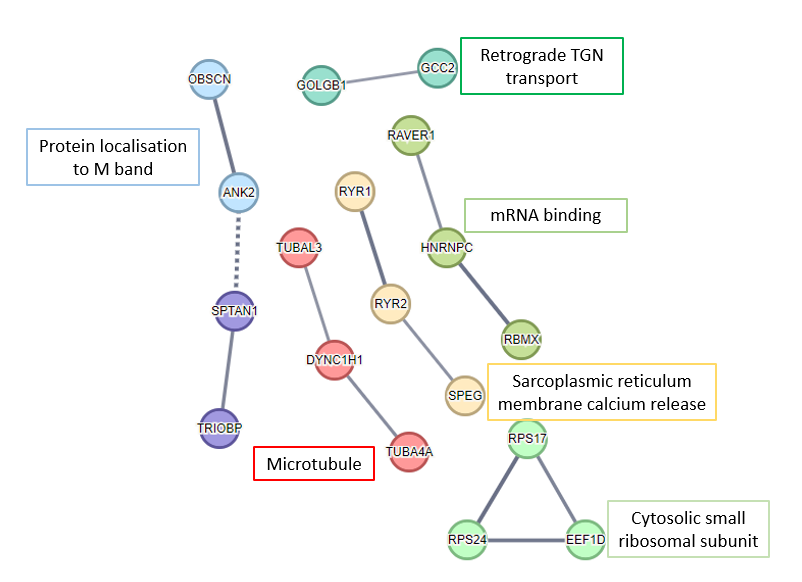

C

**Figure S4.** Gene ontology and STRING protein interaction network analysis of KMO-interacting proteins shared between KMO-Arg452 and KMO-Cys452 with confidence scores ≥40 (top ten GO terms are shown). A) GO-term enrichment analysis to derive cellular component terms associated with KMO-Arg/Cys452 interacting proteins, B) biological process terms associated with KMO-Arg/Cys452 interacting proteins, and C) STRING protein interaction network analysis to derive networks associated with KMO-Arg/Cys452 interacting proteins. Network edges show confidence where the thickness of the lines represent the strength of the supporting data.

**Table S1.** Lists of proteins with confidence scores ≥40 that specifically interacted with the KMO-Arg452 or KMO-Cys452 variants.

| **Arg452** | **Cys452** |
| --- | --- |
| HSP90AA1 | ACTA1 |
| CLASP1 | DNAH14 |
| IPO5 | HDLBP |
| HNRNPK | PLEC |
| TUBA3E | SPTBN4 |
| ZC3HAV1 | HDAC6 |
| RBBP7 | DSP |
| RPL17 | GAK |
| FSIP2 | CLASP2 |
| OTOF | EPPK1 |
| SPEN | PI4KA |
| ALDOA | CCDC88B |
| MYO9A | ATP1A2 |
| PTBP3 | ZC3H13 |
| NUMA1 | CEP135 |
| CENPE | NIN |
| MYH11 | CCDC144A |
| VAPA | AP3D1 |
| DUOX1 | CKAP5 |
| UPF2 | RPN1 |
| MTAP | SH3GLB1 |
| RYR3 | RIF1 |
| H3F3C | PPFIA2 |
| HMCN1 | TPM3 |
| SACS | PDZD2 |
| SPTBN1 | NWD1 |
| MAP1A | EIF4G3 |
| RAB15 | DARS |
| RPL28 | KRT13 |
| KIF16B | LRRFIP1 |
| PPP1R12A | CAPN1 |
| ARF3 | RPL36AL |
| ARID4A | DDX4 |
| SMG1 | ABCA10 |
| JPH2 | RBBP6 |
| MYH14 | SRP72 |
| C10orf112 | YTHDF3 |
| TIMM50 | ATP2A2 |
| PLEKHG5 | CCDC66 |
| KIF21A | HNRNPA3 |
| KIF6 | DGKA |
| GVINP1 | LIMCH1 |
| CLPB | RGPD1 |
| SOGA1 | SEPT9 |
| LRP2 | PLB1 |
| DOCK4 | CYFIP1 |
| CELSR3 | HMGB1 |
| YLPM1 | UBR4 |
| MT1G | TSC1 |
| CCDC73 | CDK12 |
| REV1 | KIAA0100 |
| SART3 | SYNJ1 |
| RABEP1 | ANKRD62 |
| GLOD4 | RGPD3 |
| ZNF831 | AFAP1L1 |
| ATRX | CCDC30 |
| PSME4 | DCTN1 |
| PPP2R2B | RNF113A |
| ACSL4 | ZNF791 |
| KIAA1551 | STAU1 |
| DHX8 | ADAM22 |
| CAMSAP2 | BCOR |
| AMPD2 | CLIP4 |
|  | TMPO |
|  | GNB2 |
|  | ZNFX1 |
|  | RPRD2 |
|  | SIPA1L3 |
|  | SVIL |
|  | IFT88 |
|  | CDC5L |
|  | WNK1 |
|  | LAMA4 |
|  | RRM2 |
|  | L1TD1 |
|  | CHD6 |
|  | ABCA13 |
|  | MAPK10 |
|  | BTN3A1 |
|  | ROCK1 |
|  | AK9 |
|  | MITF |
|  | CNTRL |
|  | RFC4 |
|  | CCDC81 |
|  | H2AFZ |
|  | GNAI3 |
|  | DOCK5 |
|  | USP1 |
|  | CDKN2A |
|  | LRRK2 |
|  | NHSL1 |
|  | EP400 |
|  | GRID2IP |
|  | CENPF |
|  | AXDND1 |
|  | ZNF571 |
|  | WDR33 |
|  | MATR3 |
|  | GRK7 |
|  | USP47 |
|  | CAPRIN2 |
|  | MYO10 |
|  | PPFIA3 |
|  | SLFN13 |
|  | PROSC |
|  | KCTD16 |

**Table S2.** GO cellular component analysis of KMO-Arg452 interacting proteins

| **GO cellular component terms** | **Number of genes in *Homo sapiens* reference list** | **Number of genes in uploaded data set that map to reference list** | **Number of genes in uploaded data set expected to map to reference list** | **Fold Enrichment of genes in data set compared to the expected number of genes** | **Raw P-value** | **False Discovery Rate** |
| --- | --- | --- | --- | --- | --- | --- |
| cortical microtubule cytoskeleton (GO:0030981) | 3 | 2 | 0.01 | 100 | 2.59E-05 | 4.68E-03 |
| microtubule minus-end (GO:0036449) | 7 | 2 | 0.02 | 96.39 | 1.80E-04 | 1.63E-02 |
| mitotic spindle midzone (GO:1990023) | 14 | 2 | 0.04 | 48.2 | 7.69E-04 | 4.63E-02 |
| microtubule end (GO:1990752) | 34 | 3 | 0.1 | 29.77 | 1.39E-04 | 1.32E-02 |
| A band (GO:0031672) | 40 | 3 | 0.12 | 25.3 | 2.26E-04 | 1.80E-02 |
| kinesin complex (GO:0005871) | 48 | 3 | 0.14 | 21.09 | 3.90E-04 | 2.98E-02 |
| myosin complex (GO:0016459) | 58 | 3 | 0.17 | 17.45 | 6.81E-04 | 4.37E-02 |
| contractile muscle fiber (GO:0043292) | 249 | 6 | 0.74 | 8.13 | 9.40E-05 | 1.25E-02 |
| sarcomere (GO:0030017) | 218 | 5 | 0.65 | 7.74 | 4.68E-04 | 3.32E-02 |
| myofibril (GO:0030016) | 239 | 5 | 0.71 | 7.06 | 7.10E-04 | 4.41E-02 |
| microtubule (GO:0005874) | 483 | 10 | 1.43 | 6.99 | 1.43E-06 | 7.09E-04 |
| supramolecular fiber (GO:0099512) | 1063 | 18 | 3.15 | 5.71 | 9.43E-10 | 9.38E-07 |
| supramolecular polymer (GO:0099081) | 1071 | 18 | 3.17 | 5.67 | 1.06E-09 | 7.05E-07 |
| supramolecular complex (GO:0099080) | 1439 | 22 | 4.27 | 5.16 | 5.11E-11 | 1.02E-07 |
| axon (GO:0030424) | 651 | 9 | 1.93 | 4.66 | 1.20E-04 | 1.25E-02 |
| polymeric cytoskeletal fiber (GO:0099513) | 818 | 11 | 2.42 | 4.54 | 2.48E-05 | 4.94E-03 |
| microtubule cytoskeleton (GO:0015630) | 1419 | 13 | 4.21 | 3.09 | 2.19E-04 | 1.82E-02 |
| neuron projection (GO:0043005) | 1336 | 12 | 3.96 | 3.03 | 4.76E-04 | 3.27E-02 |
| cytoskeleton (GO:0005856) | 2444 | 20 | 7.24 | 2.76 | 1.41E-05 | 3.12E-03 |
| extracellular exosome (GO:0070062) | 2102 | 17 | 6.23 | 2.73 | 8.95E-05 | 1.27E-02 |
| extracellular vesicle (GO:1903561) | 2131 | 17 | 6.32 | 2.69 | 1.06E-04 | 1.32E-02 |
| extracellular organelle (GO:0043230) | 2132 | 17 | 6.32 | 2.69 | 1.07E-04 | 1.25E-02 |
| extracellular membrane-bounded organelle (GO:0065010) | 2132 | 17 | 6.32 | 2.69 | 1.07E-04 | 1.18E-02 |
| cell projection (GO:0042995) | 2405 | 17 | 7.13 | 2.38 | 4.56E-04 | 3.36E-02 |
| intracellular non-membrane-bounded organelle (GO:0043232) | 5359 | 32 | 15.88 | 2.01 | 1.21E-05 | 3.43E-03 |
| non-membrane-bounded organelle (GO:0043228) | 5361 | 32 | 15.89 | 2.01 | 1.21E-05 | 3.02E-03 |
| cytosol (GO:0005829) | 5513 | 31 | 16.34 | 1.9 | 6.52E-05 | 1.08E-02 |
| intracellular organelle (GO:0043229) | 13324 | 55 | 39.49 | 1.39 | 9.96E-06 | 3.30E-03 |
| cytoplasm (GO:0005737) | 12190 | 50 | 36.13 | 1.38 | 2.07E-04 | 1.79E-02 |
| organelle (GO:0043226) | 14124 | 57 | 41.86 | 1.36 | 5.00E-06 | 1.99E-03 |
| intracellular membrane-bounded organelle (GO:0043231) | 12180 | 49 | 36.1 | 1.36 | 5.99E-04 | 3.97E-02 |
| membrane-bounded organelle (GO:0043227) | 13258 | 53 | 39.3 | 1.35 | 1.32E-04 | 1.31E-02 |
| intracellular anatomical structure (GO:0005622) | 14978 | 57 | 44.4 | 1.28 | 7.20E-05 | 1.10E-02 |

**Table S3**. KMO-Arg452-interacting proteins associated with GO terms of interest

| **cortical microtubule cytoskeleton (GO:0030981)** |  |
| --- | --- |
| **Gene name** | **Protein name** |
| NUMA1 | Nuclear mitotic apparatus protein 1 |
| CLASP1 | Isoform 3 of CLIP-associating protein 1 |
| **kinesin complex (GO:0005871)** |  |
| **Gene name** | **Protein name** |
| KIF6 | Kinesin-like protein KIF6 |
| KIF16B | Kinesin-like protein KIF16B |
| KIF21A | Kinesin-like protein KIF21A |
| **myosin complex (GO:0016459)** |  |
| **Gene name** | **Protein name** |
| MYO9A | Myosin XIA |
| MYH11 | Myosin heavy chain 11 |
| MYH14 | Myosin heavy chain 14 |
| **contractile muscle fiber (GO:0043292)** |  |
| **Gene name** | **Protein name** |
| MYH11 | Myosin heavy chain 11 |
| ALDOA | Isoform 2 of Fructose-bisphosphate aldolase A |
| RYR3 | Ryanodine receptor 3 |
| SPTBN1 | Spectrin beta chain_ non-erythrocytic 1 |
| PPP1R12A | Protein phosphatase 1 regulatory subunit 12A |
| JPH2 | Junctophilin 2 |
| **microtubule end (GO:1990752)** |  |
| **Gene name** | **Protein name** |
| CAMSAP2 | Calmodulin-regulated spectrin-associated protein 2 |
| NUMA1 | Nuclear mitotic apparatus protein 1 |
| CLASP1 | Isoform 3 of CLIP-associating protein 1 |
| **mitotic spindle midzone (GO:1990023)** |  |
| **Gene name** | **Protein name** |
| CENPE | Centromere protein E |
| NUMA1 | Nuclear mitotic apparatus protein 1 |

**Table S4.** GO biological process (A) and cellular component (B) analysis of KMO-Cys452 interacting proteins

| A) **GO biological process terms** | **Number of genes in *Homo sapiens* reference list** | **Number of genes in uploaded data set that map to reference list** | **Number of genes in uploaded data set expected to map to reference list** | **Fold Enrichment of genes in data set compared to the expected number of genes** | **Raw P-value** | **False Discovery Rate** |
| --- | --- | --- | --- | --- | --- | --- |
| centriole-centriole cohesion (GO:0010457) | 13 | 3 | 0.07 | 43.97 | 3.87E-05 | 3.90E-02 |
| regulation of synaptic vesicle recycling (GO:1903421) | 26 | 4 | 0.14 | 29.32 | 9.81E-06 | 2.97E-02 |
| synapse organization (GO:0050808) | 337 | 10 | 1.77 | 5.65 | 1.14E-05 | 2.45E-02 |
| regulation of autophagy (GO:0010506) | 363 | 10 | 1.9 | 5.25 | 2.15E-05 | 3.62E-02 |
| regulation of actin filament-based process (GO:0032970) | 382 | 10 | 2 | 4.99 | 3.32E-05 | 4.19E-02 |
| regulation of supramolecular fiber organization (GO:1902903) | 387 | 10 | 2.03 | 4.92 | 3.71E-05 | 4.01E-02 |
| supramolecular fiber organization (GO:0097435) | 596 | 14 | 3.13 | 4.48 | 2.87E-06 | 4.34E-02 |
| cell junction organization (GO:0034330) | 533 | 12 | 2.8 | 4.29 | 2.35E-05 | 3.23E-02 |
| regulation of cellular component biogenesis (GO:0044087) | 979 | 18 | 5.14 | 3.5 | 3.23E-06 | 2.44E-02 |
| positive regulation of cellular component organization (GO:0051130) | 1115 | 18 | 5.85 | 3.08 | 1.91E-05 | 3.61E-02 |
| cytoskeleton organization (GO:0007010) | 1256 | 20 | 6.59 | 3.03 | 7.50E-06 | 2.83E-02 |
| cellular response to stress (GO:0033554) | 1589 | 22 | 8.34 | 2.64 | 2.22E-05 | 3.36E-02 |
| organelle organization (GO:0006996) | 3099 | 35 | 16.26 | 2.15 | 5.05E-06 | 2.55E-02 |
| cellular component biogenesis (GO:0044085) | 2705 | 30 | 14.2 | 2.11 | 4.50E-05 | 4.25E-02 |
| cellular component organization (GO:0016043) | 5623 | 51 | 29.51 | 1.73 | 1.07E-05 | 2.70E-02 |
| cellular component organization or biogenesis (GO:0071840) | 5838 | 51 | 30.64 | 1.66 | 3.66E-05 | 4.25E-02 |

| B) **GO cellular component terms** | **Number of genes in *Homo sapiens* reference list** | **Number of genes in uploaded data set that map to reference list** | **Number of genes in uploaded data set expected to map to reference list** | **Fold Enrichment of genes in data set compared to the expected number of genes** | **Raw P-value** | **False Discovery Rate** |
| --- | --- | --- | --- | --- | --- | --- |
| centriolar subdistal appendage (GO:0120103) | 10 | 3 | 0.05 | 57.17 | 1.64E-05 | 1.48E-03 |
| hemidesmosome (GO:0030056) | 7 | 2 | 0.04 | 54.44 | 5.63E-04 | 3.20E-02 |
| ciliary transition fiber (GO:0097539) | 8 | 2 | 0.04 | 47.64 | 7.48E-04 | 3.92E-02 |
| microtubule plus-end (GO:0035371) | 24 | 3 | 0.13 | 23.82 | 2.63E-04 | 1.63E-02 |
| microtubule end (GO:1990752) | 34 | 4 | 0.18 | 22.42 | 2.95E-05 | 2.44E-03 |
| podosome (GO:0002102) | 33 | 3 | 0.17 | 17.32 | 6.84E-04 | 3.68E-02 |
| terminal bouton (GO:0043195) | 46 | 4 | 0.24 | 16.57 | 9.88E-05 | 7.02E-03 |
| cell cortex region (GO:0099738) | 35 | 3 | 0.18 | 16.33 | 8.14E-04 | 4.05E-02 |
| contractile actin filament bundle (GO:0097517) | 85 | 6 | 0.45 | 13.45 | 5.67E-06 | 7.05E-04 |
| stress fiber (GO:0001725) | 85 | 6 | 0.45 | 13.45 | 5.67E-06 | 6.64E-04 |
| actomyosin (GO:0042641) | 91 | 6 | 0.48 | 12.56 | 8.43E-06 | 8.82E-04 |
| actin filament bundle (GO:0032432) | 94 | 6 | 0.49 | 12.16 | 1.02E-05 | 1.01E-03 |
| cytoplasmic region (GO:0099568) | 308 | 9 | 1.62 | 5.57 | 3.58E-05 | 2.85E-03 |
| midbody (GO:0030496) | 206 | 6 | 1.08 | 5.55 | 7.67E-04 | 3.91E-02 |
| cell cortex (GO:0005938) | 314 | 8 | 1.65 | 4.85 | 2.52E-04 | 1.62E-02 |
| polymeric cytoskeletal fiber (GO:0099513) | 818 | 19 | 4.29 | 4.43 | 4.66E-08 | 1.85E-05 |
| centrosome (GO:0005813) | 737 | 17 | 3.87 | 4.4 | 2.86E-07 | 8.12E-05 |
| microtubule (GO:0005874) | 483 | 11 | 2.53 | 4.34 | 4.72E-05 | 3.61E-03 |
| actin cytoskeleton (GO:0015629) | 521 | 11 | 2.73 | 4.02 | 9.29E-05 | 6.84E-03 |
| cell-substrate junction (GO:0030055) | 434 | 9 | 2.28 | 3.95 | 4.71E-04 | 2.75E-02 |
| microtubule organizing center (GO:0005815) | 879 | 18 | 4.61 | 3.9 | 6.99E-07 | 1.39E-04 |
| microtubule cytoskeleton (GO:0015630) | 1419 | 26 | 7.45 | 3.49 | 1.42E-08 | 7.04E-06 |
| supramolecular fiber (GO:0099512) | 1063 | 19 | 5.58 | 3.41 | 2.51E-06 | 3.56E-04 |
| supramolecular polymer (GO:0099081) | 1071 | 19 | 5.62 | 3.38 | 2.80E-06 | 3.71E-04 |
| anchoring junction (GO:0070161) | 902 | 16 | 4.73 | 3.38 | 1.88E-05 | 1.63E-03 |
| supramolecular complex (GO:0099080) | 1439 | 25 | 7.55 | 3.31 | 8.07E-08 | 2.68E-05 |
| axon (GO:0030424) | 651 | 11 | 3.42 | 3.22 | 6.25E-04 | 3.45E-02 |
| cytoskeleton (GO:0005856) | 2444 | 41 | 12.83 | 3.2 | 2.77E-12 | 5.52E-09 |
| cell junction (GO:0030054) | 2238 | 30 | 11.74 | 2.55 | 9.06E-07 | 1.50E-04 |
| cell projection (GO:0042995) | 2405 | 31 | 12.62 | 2.46 | 1.31E-06 | 2.00E-04 |
| plasma membrane bounded cell projection (GO:0120025) | 2291 | 26 | 12.02 | 2.16 | 1.48E-04 | 9.80E-03 |
| intracellular non-membrane-bounded organelle (GO:0043232) | 5359 | 57 | 28.12 | 2.03 | 3.40E-09 | 3.38E-06 |
| non-membrane-bounded organelle (GO:0043228) | 5361 | 57 | 28.13 | 2.03 | 3.44E-09 | 2.28E-06 |
| cytosol (GO:0005829) | 5513 | 54 | 28.93 | 1.87 | 2.89E-07 | 7.19E-05 |
| protein-containing complex (GO:0032991) | 6512 | 56 | 34.17 | 1.64 | 1.56E-05 | 1.48E-03 |
| intracellular organelle (GO:0043229) | 13324 | 93 | 69.92 | 1.33 | 8.14E-07 | 1.47E-04 |
| organelle (GO:0043226) | 14124 | 97 | 74.12 | 1.31 | 3.32E-07 | 7.35E-05 |
| cytoplasm (GO:0005737) | 12190 | 82 | 63.97 | 1.28 | 3.63E-04 | 2.19E-02 |
| membrane-bounded organelle (GO:0043227) | 13258 | 88 | 69.58 | 1.26 | 1.16E-04 | 7.95E-03 |
| intracellular anatomical structure (GO:0005622) | 14978 | 98 | 78.6 | 1.25 | 6.07E-06 | 6.71E-04 |

**Table S5**. KMO-Cys452-interacting proteins associated with GO terms of interest

| **centriole-centriole cohesion (GO:0010457)** |  |
| --- | --- |
| **Gene name** | **Protein name** |
| NIN | Isoform 3 of Ninein |
| CEP135 | Centrosomal protein 135 |
| DCTN1 | Dynactin subunit 1 |
| **regulation of synaptic vesicle recycling (GO:1903421)** |  |
| **Gene name** | **Protein name** |
| CYFIP1 | Cytoplasmic FMR1-interacting protein 1 |
| ABCA13 | ATP binding cassette subfamily A member 13 |
| ROCK1 | Rho-associated protein kinase 1 |
| LRRK2 | Leucine-rich repeat serine/threonine-protein kinase 2 |
| **synapse organization (GO:0050808)** |  |
| **Gene name** | **Protein name** |
| SPTBN4 | Spectrin beta, non-erythrocytic 4 |
| CLASP2 | CLIP-associating protein 2 |
| PPFIA2 | PTPRF interacting protein alpha 2/ Liprin alpha 2 |
| TSC1 | TSC complex subunit 1 |
| CYFIP1 | Cytoplasmic FMR1-interacting protein 1 |
| STAU1 | Staufen double-stranded RNA binding protein 1 |
| LRRK2 | Leucine-rich repeat serine/threonine-protein kinase 2 |
| PPFIA3 | PTPRF interacting protein alpha 3 |
| DCTN1 | Dynactin subunit 1 |
| HDAC6 | Isoform 2 of Histone deacetylase 6 |
| **terminal bouton (GO:0043195)** |  |
| **Gene name** | **Protein name** |
| AP3D1 | AP-3 complex subunit delta-1 |
| CYFIP1 | Cytoplasmic FMR1-interacting protein 1 |
| SYNJ1 | Synaptojanin-1 |
| LRRK2 | Leucine-rich repeat serine/threonine-protein kinase 2 |
| **regulation of autophagy (GO:0010506)** |  |
| **Gene name** | **Protein name** |
| UBR4 | Ubiquitin protein ligase E3 component n-recognin 4 |
| CAPN1 | Calpain 1 |
| TSC1 | TSC complex subunit 1 |
| WNK1 | WNK lysine deficient protein kinase 1 |
| ROCK1 | Rho-associated protein kinase 1 |
| LRRK2 | Leucine-rich repeat serine/threonine-protein kinase 2 |
| HDAC6 | Isoform 2 of Histone deacetylase 6 |
| SH3GLB1 | SH3 domain containing GRB2 like, endophilin B1 |
| HMGB1 | High mobility group box 1 |
| GNAI3 | G protein subunit alpha i3 |
| **supramolecular fiber organization (GO:0097435)** |  |
| **Gene name** | **Protein name** |
| PLEC | Isoform 2 of Plectin |
| CCDC88B | Coiled-coil domain containing 88B |
| CKAP5 | Cytoskeleton-associated protein 5 |
| CLASP2 | CLIP-associating protein 2 |
| DSP | Desomoplakin |
| NIN | Isoform 3 of Ninein |
| EPPK1 | Epiplakin 1 |
| TPM3 | Isoform 2 of Tropomyosin alpha-3 chain |
| CLIP4 | CAP-Gly domain-containing linker protein 4 |
| KRT13 | Keratin_ type I cytoskeletal 13 |
| ACTA1 | Actin_ alpha skeletal muscle |
| SVIL | Supervillin |
| HDAC6 | Isoform 2 of Histone deacetylase 6 |
| CDKN2A | Cyclin dependent kinase inhibitor 2A |
| **microtubule plus-end (GO:0035371)** |  |
| **Gene name** | **Protein name** |
| CKAP5 | Cytoskeleton-associated protein 5 |
| DCTN1 | Dynactin subunit 1 |
| CLIP4 | CAP-Gly domain-containing linker protein 4 |
| **centriolar subdistal appendage (GO:0120103)** |  |
| **Gene name** | **Protein name** |
| NIN | Isoform 3 of Ninein |
| CNTRL | Centriolin |
| DCTN1 | Dynactin subunit 1 |
| **contractile actin filament bundle (GO:0097517)** |  |
| **Gene name** | **Protein name** |
| TPM3 | Isoform 2 of Tropomyosin alpha-3 chain |
| LIMCH1 | LIM and calponin homology domains 1 |
| SIPA1L3 | Signal induced proliferation associated 1 like 3 |
| AFAP1L1 | Actin filament associated protein 1 like 1 |
| ACTA1 | Actin_ alpha skeletal muscle |
| SEPT9 | Septin 9 |

**Table S6.** Proteins with confidence scores ≥40 for both KMO variants and that interacted with both KMO-Arg452 and KMO-Cys452.

**Both Arg452 and Cys452**

SYNE1

KMO

DYNC1H1

TUBA4A

AHNAK

GOLGB1

OBSCN

FAM179B

MYH10

RBMX

RAVER1

GSR

TTK

RYR2

KRT17

LARP1B

BRWD1

MAP4

RPLP0P6

RCN2

CEP290

PABPC1L

LDHA

GCC2

ATP1A3

BAX

SPTAN1

BAG6

ERC1

RPS24

SUPT6H

TUBAL3

RGPD5

SAMHD1

EEF1D

TRIOBP

RPS17

RYR1

ANK2

HELLS

HNRNPC

EIF4B

FGD2

HAX1

SPEG

GTF2IRD2

**Table S7.** GO biological process (A) and cellular component (B) analysis of KMO-Arg/Cys452 interacting proteins

| **A) GO biological process terms** | **Number of genes in *Homo sapiens* reference list** | **Number of genes in uploaded data set that map to reference list** | **Number of genes in uploaded data set expected to map to reference list** | **Fold Enrichment of genes in data set compared to the expected number of genes** | **Raw P-value** | **False Discovery Rate** |
| --- | --- | --- | --- | --- | --- | --- |
| protein localization to M-band (GO:0036309) | 2 | 2 | 0 | 100 | 4.89E-06 | 1.06E-02 |
| regulation of SA node cell action potential (GO:0098907) | 3 | 2 | 0.01 | 100 | 1.46E-05 | 2.77E-02 |
| regulation of atrial cardiac muscle cell action potential (GO:0098910) | 4 | 2 | 0.01 | 100 | 2.92E-05 | 4.91E-02 |
| sarcoplasmic reticulum calcium ion transport (GO:0070296) | 15 | 3 | 0.03 | 89.48 | 4.67E-06 | 1.18E-02 |
| protein localization to organelle (GO:0033365) | 739 | 10 | 1.65 | 6.05 | 4.24E-06 | 1.28E-02 |
| cytoskeleton organization (GO:0007010) | 1256 | 15 | 2.81 | 5.34 | 4.75E-08 | 3.59E-04 |
| organelle organization (GO:0006996) | 3099 | 21 | 6.93 | 3.03 | 7.64E-07 | 2.89E-03 |
| cellular component organization or biogenesis (GO:0071840) | 5838 | 32 | 13.05 | 2.45 | 7.98E-09 | 1.21E-04 |
| cellular component organization (GO:0016043) | 5623 | 30 | 12.57 | 2.39 | 8.92E-08 | 4.49E-04 |

| **B) GO cellular component terms** | **Number of genes in *Homo sapiens* reference list** | **Number of genes in uploaded data set that map to reference list** | **Number of genes in uploaded data set expected to map to reference list** | **Fold Enrichment of genes in data set compared to the expected number of genes** | **Raw P-value** | **False Discovery Rate** |
| --- | --- | --- | --- | --- | --- | --- |
| junctional sarcoplasmic reticulum membrane (GO:0014701) | 9 | 2 | 0.02 | 99.42 | 1.74E-04 | 1.65E-02 |
| costamere (GO:0043034) | 18 | 2 | 0.04 | 49.71 | 7.31E-04 | 4.69E-02 |
| sarcoplasmic reticulum (GO:0016529) | 77 | 3 | 0.17 | 17.43 | 6.81E-04 | 4.51E-02 |
| sarcolemma (GO:0042383) | 142 | 5 | 0.32 | 15.75 | 1.59E-05 | 2.64E-03 |
| Z disc (GO:0030018) | 130 | 4 | 0.29 | 13.77 | 2.02E-04 | 1.83E-02 |
| I band (GO:0031674) | 144 | 4 | 0.32 | 12.43 | 2.99E-04 | 2.47E-02 |
| myofibril (GO:0030016) | 239 | 6 | 0.53 | 11.23 | 1.46E-05 | 2.65E-03 |
| contractile muscle fiber (GO:0043292) | 249 | 6 | 0.56 | 10.78 | 1.84E-05 | 2.82E-03 |
| sarcomere (GO:0030017) | 218 | 5 | 0.49 | 10.26 | 1.23E-04 | 1.43E-02 |
| cell cortex (GO:0005938) | 314 | 6 | 0.7 | 8.55 | 6.74E-05 | 8.38E-03 |
| cytoplasmic region (GO:0099568) | 308 | 5 | 0.69 | 7.26 | 6.03E-04 | 4.28E-02 |
| supramolecular fiber (GO:0099512) | 1063 | 13 | 2.38 | 5.47 | 3.58E-07 | 1.42E-04 |
| supramolecular polymer (GO:0099081) | 1071 | 13 | 2.39 | 5.43 | 3.90E-07 | 1.29E-04 |
| supramolecular complex (GO:0099080) | 1439 | 17 | 3.22 | 5.29 | 5.18E-09 | 3.44E-06 |
| cytoskeleton (GO:0005856) | 2444 | 19 | 5.46 | 3.48 | 4.19E-07 | 1.19E-04 |
| extracellular vesicle (GO:1903561) | 2131 | 14 | 4.76 | 2.94 | 1.52E-04 | 1.68E-02 |
| extracellular organelle (GO:0043230) | 2132 | 14 | 4.77 | 2.94 | 1.53E-04 | 1.60E-02 |
| extracellular membrane-bounded organelle (GO:0065010) | 2132 | 14 | 4.77 | 2.94 | 1.53E-04 | 1.52E-02 |
| extracellular exosome (GO:0070062) | 2102 | 13 | 4.7 | 2.77 | 5.08E-04 | 4.05E-02 |
| intracellular non-membrane-bounded organelle (GO:0043232) | 5359 | 32 | 11.98 | 2.67 | 7.83E-10 | 1.56E-06 |
| non-membrane-bounded organelle (GO:0043228) | 5361 | 32 | 11.98 | 2.67 | 7.91E-10 | 7.87E-07 |
| cell projection (GO:0042995) | 2405 | 14 | 5.38 | 2.6 | 5.39E-04 | 4.13E-02 |
| cytosol (GO:0005829) | 5513 | 28 | 12.32 | 2.27 | 1.81E-06 | 3.60E-04 |
| protein-containing complex (GO:0032991) | 6512 | 26 | 14.56 | 1.79 | 6.45E-04 | 4.43E-02 |
| nucleus (GO:0005634) | 7645 | 29 | 17.09 | 1.7 | 5.92E-04 | 4.36E-02 |
| intracellular organelle (GO:0043229) | 13324 | 45 | 29.78 | 1.51 | 7.08E-08 | 3.52E-05 |
| cytoplasm (GO:0005737) | 12190 | 40 | 27.25 | 1.47 | 6.28E-05 | 8.33E-03 |
| intracellular membrane-bounded organelle (GO:0043231) | 12180 | 39 | 27.22 | 1.43 | 2.46E-04 | 2.13E-02 |
| organelle (GO:0043226) | 14124 | 45 | 31.57 | 1.43 | 1.20E-06 | 2.65E-04 |
| membrane-bounded organelle (GO:0043227) | 13258 | 42 | 29.63 | 1.42 | 4.04E-05 | 5.74E-03 |
| intracellular anatomical structure (GO:0005622) | 14978 | 46 | 33.48 | 1.37 | 8.35E-07 | 2.08E-04 |

|  | **Controls (n=170)** | **SZ (n=156)** | **Test statistic** | **p-value** |
| --- | --- | --- | --- | --- |
| **Age (years)** | 37.6 ± 15.4 | 36.0 ± 13.5 | t=0.97 | .33 |
| **Sex (M/F)** | 94/76 | 104/52 | Χ^2^=4.41 | .036 |
| **rs1053230 genotype (CC/CT/TT)** | 137/33/0 | 111/40/5 | Χ^2^=7.81 | .020 |
| **Digit symbol coding task score** | 75.1 ± 17.6 | 59.1 ± 19.1 | t=7.76 | <.001 |
| **Digit sequencing task score** | 20.9 ± 4.4 | 17.5 ± 5.0 | t=6.30 | <.001 |
| **Antipsychotic type, n(%):**  **First generation**  **Second generation**  **Multiple**  **None** |  | 119 (76.3)  11 (7.1)  19 (12.2)  7 (4.5) |  |  |

**Table S8.** Characteristics of the cohort used for association of rs1053230 genotype with clinical parameters related to schizophrenia
